# High oil accumulation in tuber of yellow nutsedge compared to purple nutsedge is associated with more abundant expression of genes involved in fatty acid synthesis and triacylglycerol storage

**DOI:** 10.1101/2020.10.25.325241

**Authors:** Hongying Ji, Dantong Liu, Zhenle Yang

## Abstract

Yellow nutsedge is a specific plant species that contains significant amounts of both starch and oil as the main reserves in storage tuber. Its tuber can accumulate up to 35% oil of dry weight, perhaps the highest level observed in the tuber tissues of plant kingdom. To gain insight into the molecular mechanism that leads to high oil accumulation in yellow nutsedge, gene expression profiles of oil production pathways involved carbon metabolism, fatty acid synthesis, triacylglycerol synthesis, and triacylglycerol storage during tuber development were compared with purple nutsedge, a very close relative of yellow nutsedge that is poor in oil accumulation. Compared with purple nutsedge, the high oil content in yellow nutsedge was associated with much higher transcripts for seed-like oil-body proteins, almost all fatty acid synthesis enzymes, and specific key enzymes of plastid Rubisco bypass as well as malate and pyruvate metabolism. However, transcript levels for carbon metabolism toward pyruvate generation were comparable and for triacylglycerol synthesis were similar in both species. Two seed-like master transcription factors ABI3 and WRI1 were found to display similar temporal transcript patterns but be expressed at 6.5- and 14.3-fold higher levels in yellow nutsedge than in purple nutsedge, respectively. A weighted gene co-expression network analysis revealed that *ABI3* is in strong transcriptional coordination with *WRI1* and other key oil-related genes. Together, these results implied that plastidial pyruvate availability and fatty acid synthesis, along with triacylglycerol storage in oil body, rather than triacylglycerol synthesis in endoplasmic reticulum, are the major factors responsible for high oil production in tuber of yellow nutsedge, and ABI3 is most likely to play a critical role in regulating oil accumulation. This study will aid understanding in underlying molecular mechanism controlling carbon partitioning toward oil production in oil-rich tuber and provide valuable reference for enhancing oil accumulation in non-seed tissues of crops through genetic breeding or metabolic engineering.

## Introduction

Yellow nutsedge (*Cyperus esculentus* L.) is a grass-like C_4_ plant species of the sedge family (Cyperaceae). Its tuber can be edible raw, dried, cooked, and roasted, or consumed in wheat-like flour and milky beverage forms (Sanchez-Zapata et al, 2012). As an unconventional underground tuber crop, yellow nutsedge is quite unique since it is the only one plant species so far that can accumulate oil at high levels (up to 35% of dry weight) (Codina-Torrella et al, 2015) in its tuber tissues, in striking contrast to common root/tuber crops such as potato, sweet potato, cassava, and sugar beet that have very low levels of oil while store exclusively starch or sugars as the major reserves in their storage organs. Compared to conventional oilseed crops including soybean, rapeseed, and peanut, yellow nutsedge is endowed with many advantage characteristics such as wide soil adaptability, strong fertility, short growth period, little maintenance, rare disease and pest harm, and high tuber yield (up to 12 t·ha^-1^) (Makareviciene et al, 2013; Liu et al, 2020). Even so, yellow nutsedge is still an underutilized and nonpopular oil crop around the world, most possibly due to be particularly laborious in cultivating and harvesting such a crop. Moreover, the mechanism of oil accumulation in yellow nutsedge is largely unknown at the molecular level, which hinders its potential application and development in oil production.

Nevertheless, yellow nutsedge is expected to be a promising candidate and new alternative for plant oil production from non-seed tissues to meet the worldwide growing demand of vegetable oil consumption. Given significant amounts of all the major storage components, i.e., starch, oil, sugar, and protein, are produced in its tubers (Arafat et al, 2009; Sanchez-Zapata et al, 2012; Codina-Torrella et al, 2015), yellow nutsedge is regarded as a novel model system to study carbon flux toward oil synthesis in underground storage tissues. Biochemical analysis showed that the accumulation patterns of oil, starch and soluble sugars appeared in a sequential way in developing tuber (Turesson et al, 2010), where sugar loading is paralleled with the beginning of starch accumulation upon the initialization of tuber development, and followed by later occurrence of oil accumulation accompanied by the significantly deceased sugar levels. However, it is still unclear how carbon is partitioned and how oil synthesis is directed and regulated in tuber. Currently, the scarcity of molecular resources and genomic information available for this crop also hinders our understanding of the underlying molecular mechanism governing oil production.

The synthesis of plant oil in the form of triacylglycerol (TAG) is well understood in oil-rich seed plants particularly in model plant Arabidopsis (Bates et al, 2013). In general, oil synthesis mainly involves *de novo* synthesis of fatty acids in plastid and their sequential acylation of the glycerol backbone via acyl-CoA-dependent and - independent reactions in endoplasmic reticulum (ER). Although oil synthesis in oilseed plants is well characterized and the core metabolic processes or pathways are considered to be conserved among diverse plants, the biochemical and molecular details occurred in underground oil-rich sink organs such as yellow nutsedge tuber remain largely unknown.

It is noted that a closely related species of yellow nutsedge, purple nutsedge (*Cyperus rotundus*), contains less than 5% oil of dry weight in its tuber (Asenjo 1941; Hauser 1963; Duble 1970; Stoller and Weber 1974), while stores high amounts of carbohydrates (starch and sugar). Moreover, the habit and habitat as well the growth and development of the two species are quite similar (Williams, 1982). Thus, purple nutsedge provides a good reference for exploring the biological mechanism of why and how tuber accumulate so high contents of oil in yellow nutsedge. To decipher the molecular basis controlling such considerable differences in oil content, and better understand the regulation mechanisms of oil accumulation in oil-rich tuber of yellow nutsedge, we made a comparative analysis in this study of the global gene expressions related to oil accumulation in developing tubers of yellow nutsedge with purple nutsedge. Our results revealed that high oil accumulation in tuber of yellow nutsedge is tightly associated with more abundant transcripts for fatty acid synthesis and triacylglycerol storage as compared to purple nutsedge.

## Materials and methods

### Plant materials and growth conditions

The two plant species used in this study are the cultivated varieties of *Cyperus esculentus* L. var *sativus* Boeck and *Cyperus rotundus L*., respectively. Undamaged and healthy tubers harvested last year were chosen as seed materials. Before sowing, seed tubers were kept at 4°C for 4 days and then wrapped in moist blotting paper for 2 d to promote maximum viability and uniform sprouting. After that, tubers were sown in sterilized soil mixture (commercial nutrient soil: vermiculite, 1:1) at 1.0-2.0 cm depth in 22×21 cm (diameter×height) plastic pots. Planting density was one tuber per pot. Pots with seed tubers were placed on an experimental plot without any other plants nearby on the condition of natural light and temperature at a location of 116°13’18”E, 39°59’32”N in Beijing, China. Seed tubers were allowed to sprout in moist soil and new plants were grown until maturity. All plants were fertilized biweekly with liquid nutrient Murashige and Skoog (MS) solution (Murashige and Skoog, 1962) and were watered as necessary to keep the soil appropriate moist throughout plant growth period. When needed, plant leaves were clipped after shoot emergence to maintain a height of no more than 50 cm.

The whole plant growth period experienced from mid-April to mid-September of 2016, with climate feature on the average being: April, 10.7 h night at 10°C /13.3 h day at 22°C; May, 9.6 h night at 13°C /14.4 h day at 26°C; June, 9 h night at 18°C /15 h day at 31°C; July, 9.3 h night at 21°C /14.7 h day at 32°C; August, 10.3 h night at 20°C /13.7 h day at 30°C; September, 11.5 h night at 14°C /12.5 h day at 26°C. Humidity during these months is in the range of 32-67%.

Fresh new tubers at different developmental stages were randomly selected and harvested, and then immediately washed and cleaned free of soil and fibrous scaly appendages, and stored in −80°C before use in subsequent experiments.

### Quantitative analyses of proximate components in tubers

Before analysis, tubers were crushed in liquid N2 in a mortar.

Oil content and the fatty acid composition were determined by using the method as described previously (Liu et al, 2020).

Proteins were extracted from tuber samples by homogenizing in ice cold buffer (0.5 mol/L NaCl, 0.001 mol/L EDTA, 1% (w/v) SDS, and 0.02 mol/L Tris-HCl, pH 7.5) and was centrifuged at 15,000 g at 4°C for 30 min. Protein in the supernatant was measured by a BCA Protein Assay (Thermo Fisher (China) Scientific Inc., Ltd).

For determination of starch and sugar, powered tuber samples were homogenized in 80% (v/v) ethanol for 10 min and then centrifuged at 6,000 g for 10 min. The liquid supernatant was used for sugar analysis while the remaining sediment for starch analysis. Soluble sugars were quantified according to the procedure used before (Yang et al, 2018). Starch was measured via the method of two-wavelength iodine binding colorimetry (Zhu et al, 2008).

All the determinations were performed in at least triplicates.

### RNA isolation

Total RNA was extracted according to a protocol (Yang et al., 2016) modified from CTAB-based method (Gambino et al, 2008). Pure high-quality RNA samples dissolved in non-RNase water were stored in liquid N2 for further analysis.

### RNA deep sequencing

Before sequencing, the purity, concentration and integrity of RNA samples were assessed using Nanodrop 2000 spectrophotometer, Qubit 2.0 Fluorometer, and Agilent 2100 Bioanalyzer to ensure samples qualified for RNA sequencing. RNA sequencing was carried out on an Illumina Hiseq4000 sequencing and analysis platform (Biomarker Technologies Corporation, China). High-quality reads filtered from generated raw reads (raw data) were *de novo* assembled by using the Trinity program (Grabherr et al, 2011). Annotation for assembly sequences (> 200 bp) was conducted using a BLAST homology search against the NCBI NR database (ftp://ftp.ncbi.nlm.nih.gov/blast/db/), Swiss-Prot (http://www.ebi.ac.uk/uniprot/), GO (http://www.geneontology.org/), COG (http://www.ncbi.nlm.nih.gov/COG/), KOG (ftp://ftp.ncbi.nih.gov/pub/COG/KOG/), eggnog (http://eggnog.embl.de) and KEGG (http://www.genome.jp/kegg/). Transcript levels of genes were quantified based on the number of fragments per kilobase of exon model per million mapped reads (FPKM) (Mortazavi et al, 2008).

### Gene co-expression network construction

Co-expression network was constructed based on the transcript data sets using the method of weighted gene co-expression network analysis (WGCNA) (Zhang and Horvath, 2005; Langfelder and Horvath, 2008). Pearson’s correlation coefficient above a threshold of 0.9 for a pair of gene expression was used to filter the gene pairwise connection. The resulting network was displayed with Cytoscape software platform (Shannon et al, 2003; www.cytoscape.org).

## Results

### Tuber Oil Contents Differed Markedly Between Yellow and Purple Nutsedge

Under our experimental conditions, new shoots from seed tubers of the two nutsedges appeared above soil at 5-9 days after sowing on April 16, 2016. New tubers began to present from 6 to 8 weeks after shoot emergence. Both species lasted around five months for their growth and development.

Analyses of the proximate constituents of mature tubers (Fig. 1A) indicated that there are marked differences between two species in the contents of major storage reserves including starch, oil, sugar, and protein on a tuber dry weight basis (Fig. 1B), where starch was the component at highest level in the tubers. The most notable difference is that yellow nutsedge stored more than 25% oil of dry weight while purple nutsedge contained less than 3% oil in mature tuber, indicating that there is around 10-fold difference in oil content.

**Fig. 1.**
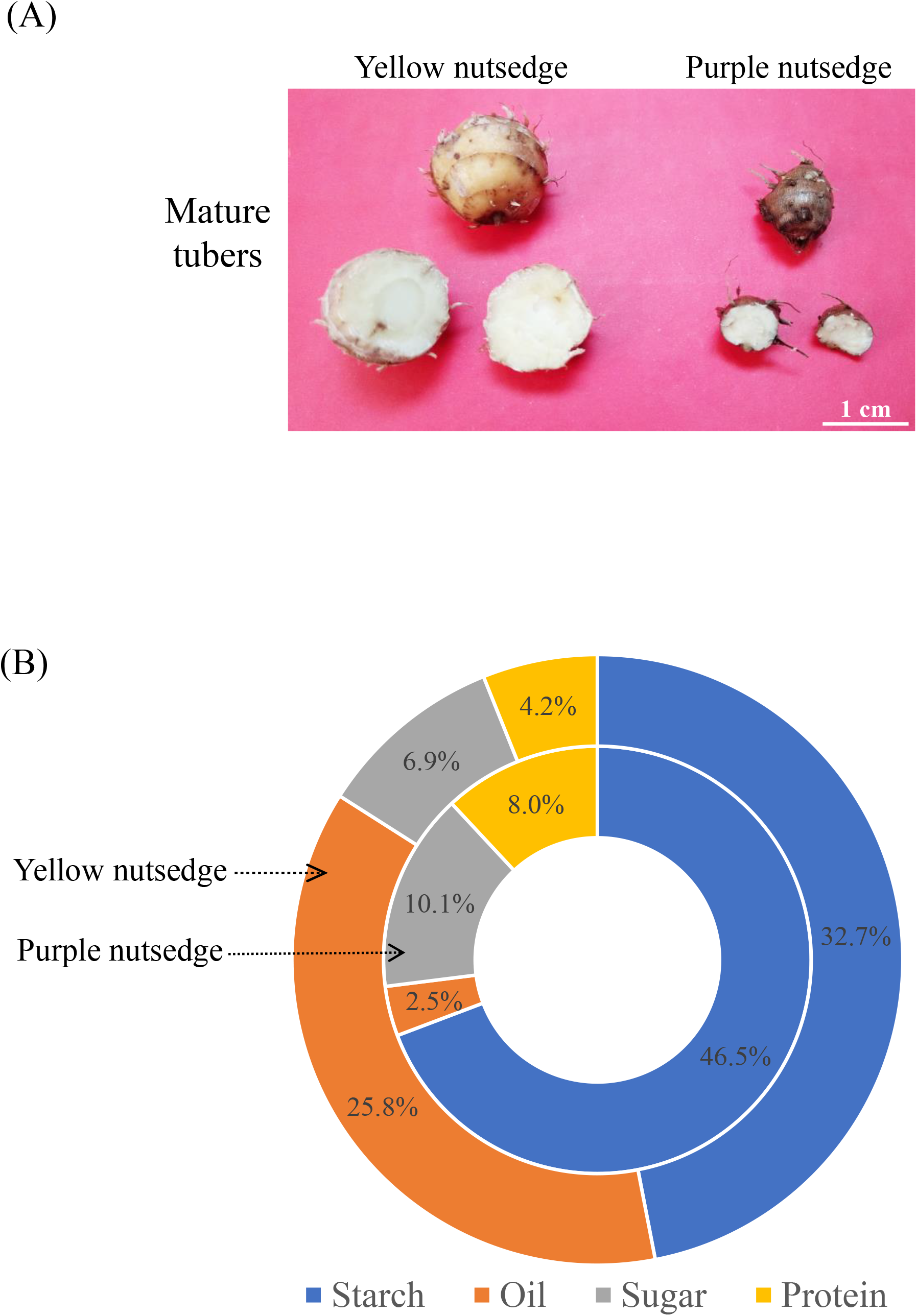
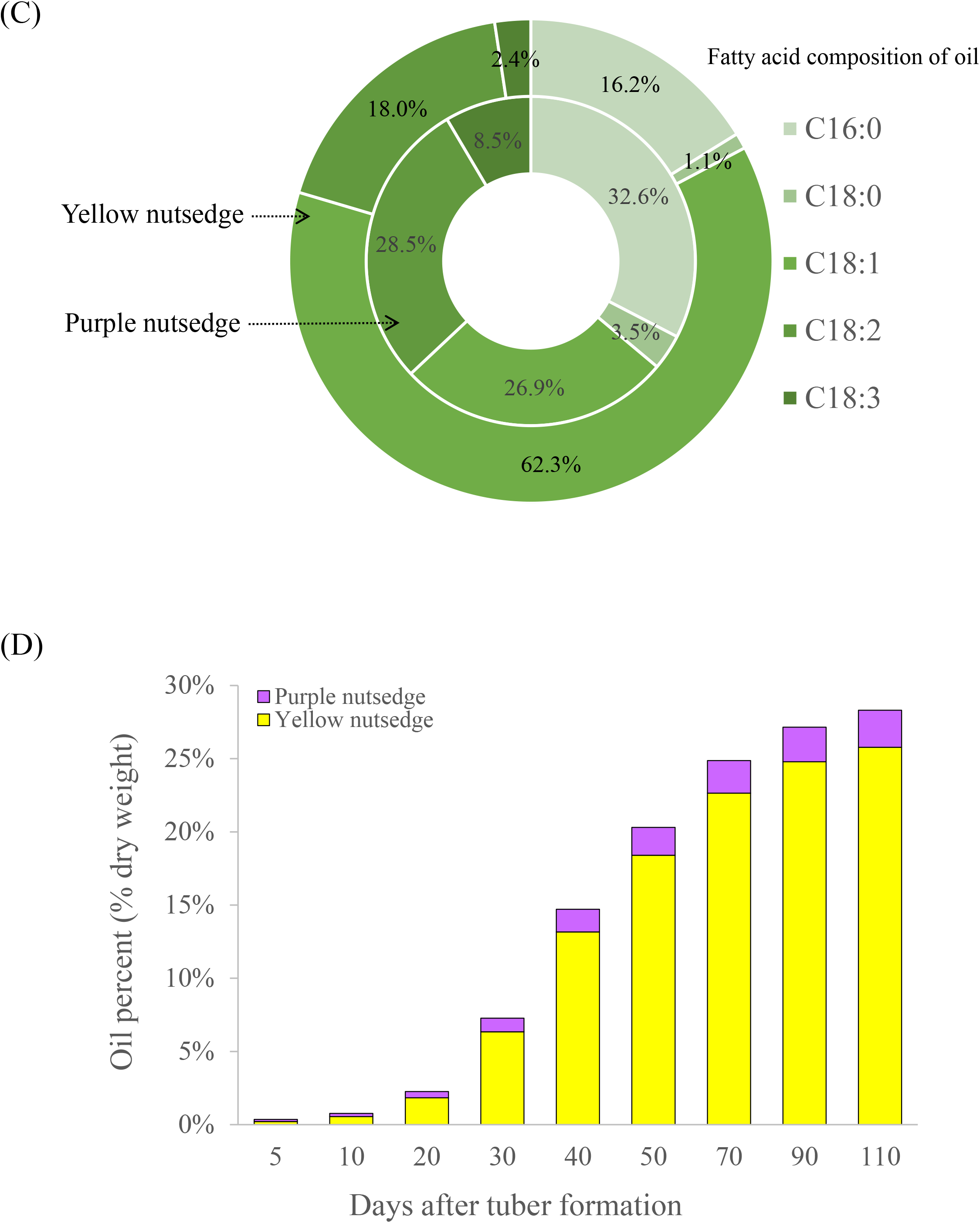
Proximate components of tubers from yellow and purple nutsedge. (A) Open tubers. (B) Proximate composition (as % dry weight) of mature tubers. (C) Fatty acid composition of oil in mature tubers. (D) Percentage of oil content on basis of tuber dry weight during tuber development. Values represent means±SD (*N*= 3).

Analysis of fatty acid composition of oil from mature tubers showed that yellow nutsedge was predominated with oleic acid (C18:1) that accounted for more than 60% of total fatty acids, while purple nutsedge was presented with palmitic acid (16:0), C18:1, and linoleic acid (18:2) as major fatty acids, with concentrations ranging from 25% to 35% (Fig. 1C). In yellow nutsedge tuber, C16:0 and C18:2 were the second most, but each of them constituted less than 20% of total fatty acids. All these results indicated that significant difference also occurred in fatty acid composition of tuber oil between these two species, where purple nutsedge contained less oleic acid and more saturated fatty acids than yellow nutsedge.

To check whether there is also a difference of oil accumulation occurred in developing tubers, the changes in oil contents during tuber development were determined for two species. The oil accumulation patterns in the development period spanning around 110 days after tuber formation (DAF) was shown in Fig. 1D. The results showed that the oil accumulation in the two types of tubers continued to increase throughout the tuber development. In all developmental stages, however, yellow nutsedge tubers contained a significantly higher percentage of oil than the purple nutsedge tubers.

On a per dry tuber basis, both nutsedges had relatively slow oil accumulation at early stage (0-20 DAF) and late stage (50-110 DAF), much lower than that of middle stage (20-50 DAF) where oil was rapidly produced, suggesting that oil accumulation was directly related to the tuber development. The rate of oil accumulation of yellow nutsedge tubers in each of three development stages was significantly greater than that of purple nutsedge tubers, in which the accumulation rates with 0.11% oil·d^-1^, 0.56% oil·d^-1^, and 0.12% oil·d^-1^ at earl-, mid- and later-stage, respectively, for yellow nutsedge is in sharp contrast to the corresponding values with 0.02% oil·d^-1^, 0.05% oil·d^-1^, and 0.01% oil·d^-1^ for purple nutsedge.

Overall, the striking differences in oil content and the fatty acid composition were present between these two types of tubers, suggesting that there exists distinct transcriptional control of oil production for them.

### Overall Level of Transcripts for Oil Production Is Higher in Yellow Nutsedge Than in Purple Nutsedge

To uncover the difference in the transcriptional control of oil production between the two species, we systematically conducted comparative transcriptome analyses of oil-related genes in developing tubers. Tuber samples at three different developmental stages (i.e. the early stage 20 DAF, the middle stage 50 DAF and the late stage 90 DAF) were used for comparative transcript analysis.

Oil production involves the conversion of sucrose up to TAG assembly or storage, primarily including cytosolic and plastidial carbon metabolism toward pyruvate generation in plastid, FA synthesis in plastid, TAG synthesis in ER and TAG storage in oil body or lipid droplet. These four major metabolic processes are related to expression of more than 400 genes (Supplemental Table S2 and S3). In this study, transcript levels were represented by FPKM (fragments per kilobase of exon model per million mapped reads) per protein, where multiple transcripts for genes encoding for isoforms or subunits of the same protein family were summed. Among these metabolic pathways, more than 70% of the transcripts was associated with genes involved in TAG storage and only 8% and 3% with those in FA synthesis and TAG synthesis, respectively, in yellow nutsedge tuber (Fig. 2A). In contrast, the transcripts related to carbon metabolism were most abundant while those of TAG storage were the least in purple nutsedge tuber.

**Fig. 2.**
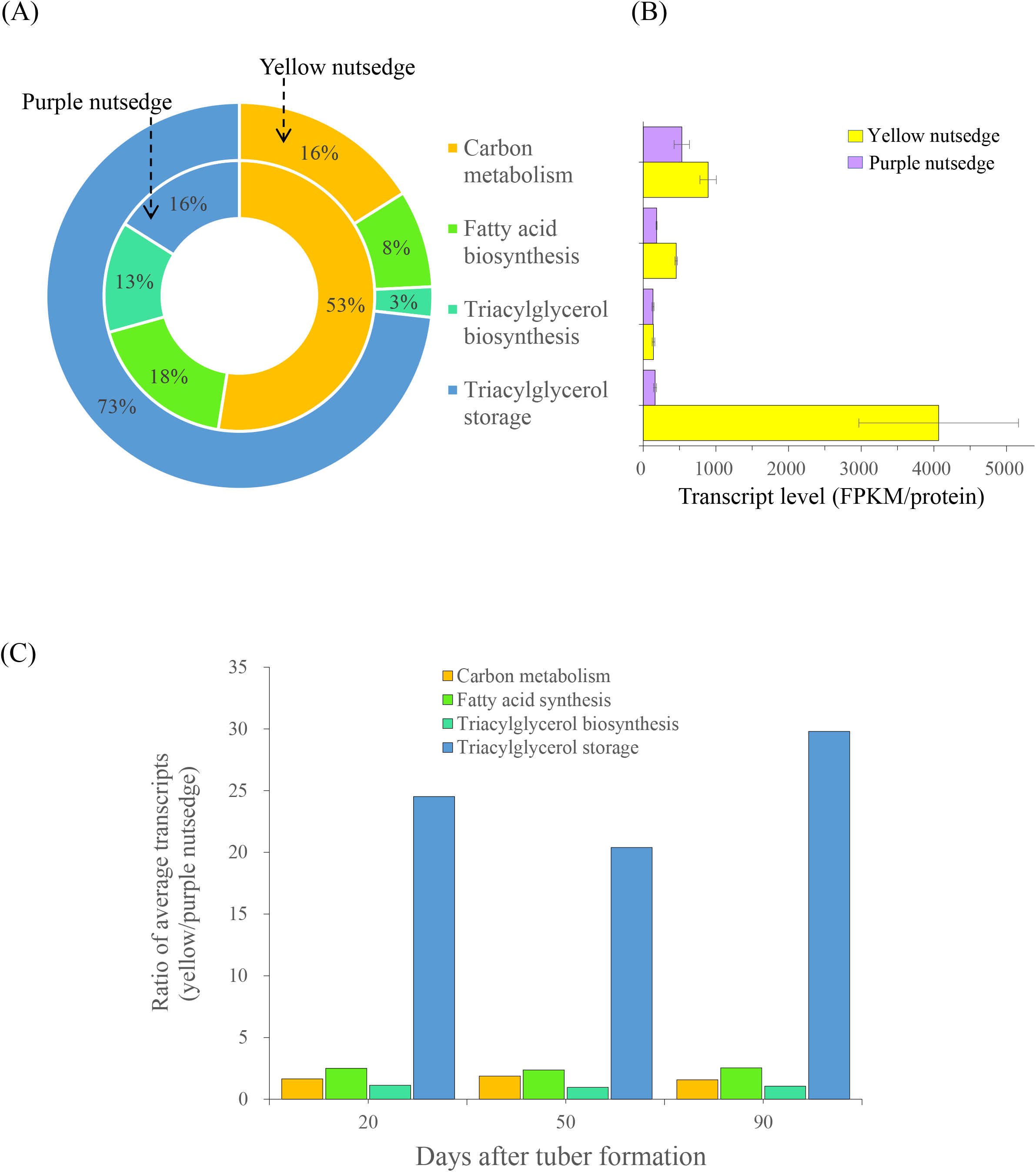
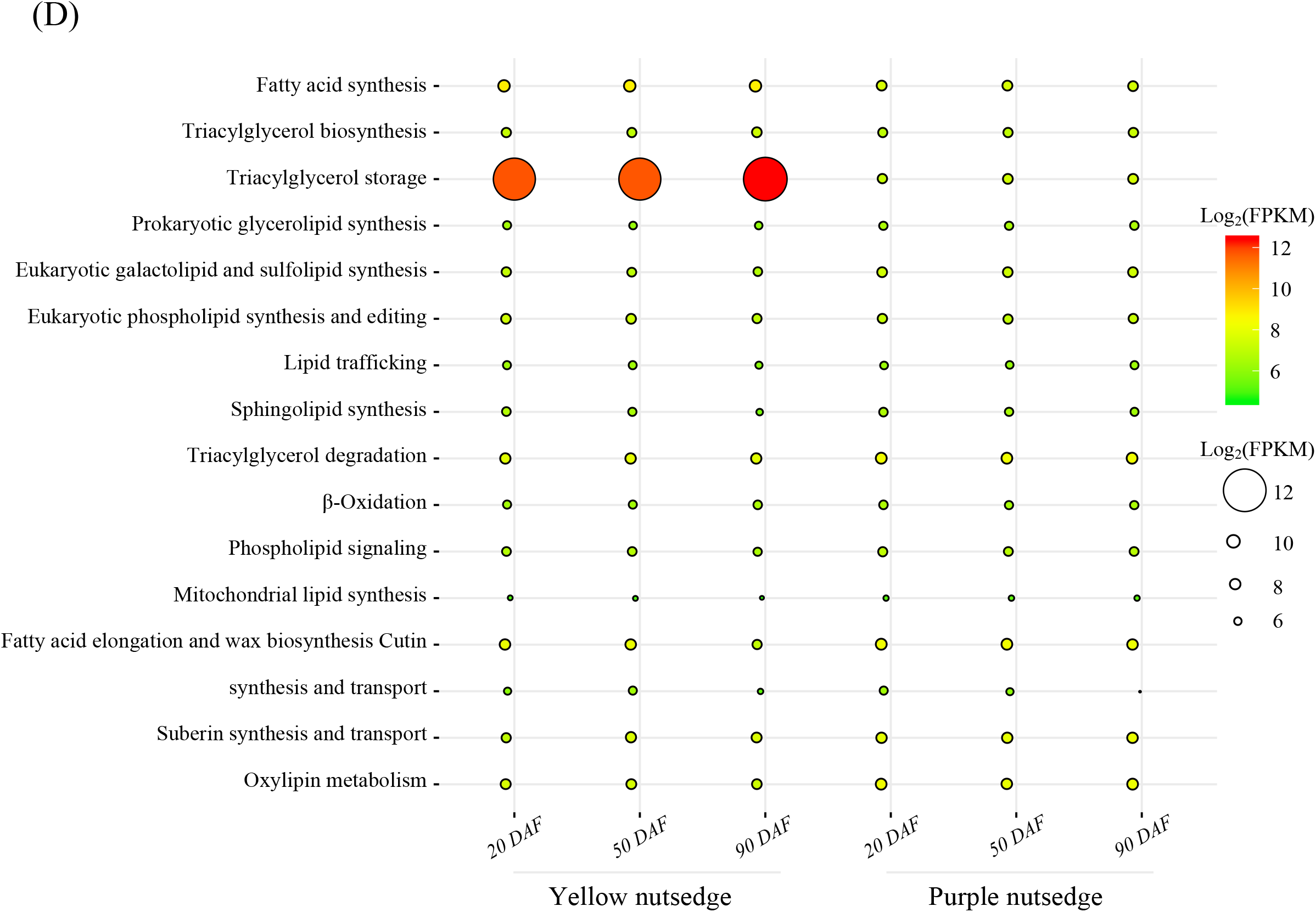
Expression pattern for the selected four pathways related to oil production in tubers. (A) The relative distribution of transcripts among the four metabolic pathways from sucrose to TAG storage. (B) Transcript level (FPKM/protein) of each pathway in two species. The FPKM values for subunits of a protein and for multiple isoforms were summed. The data are averaged on all developing stages of tubers with error bars indicating their standard deviation. (C) Ratio of transcriptional levels between two species at three development stages. (D) Temporal transcripts for lipid-related metabolic pathways during tuber development. DAF, days after tuber formation.

For every metabolic pathway, transcript levels were on average higher in yellow nutsedge than in purple nutsedge (Fig. 2B). The largest difference was noted for TAG storage, for which the transcript level was more than 25-fold higher in yellow nutsedge compared to purple nutsedge. Clear difference was also present for FA synthesis, where there is 2.5 times higher in yellow nutsedge than in purple nutsedge. Unexpectedly, there is no substantial difference of carbon metabolism and TAG synthesis between two species. A similar contrast between the two plants also occurred across the three developmental stages of tubers (Fig. 2C).

Notably, transcript patterns for other lipid-related metabolic pathways were comparable in yellow and purple nutsedge (Fig. 2D).

Overall, the results mentioned above suggest that transcriptional control of genes involved in TAG storage along with FA synthesis rather than TAG synthesis might be the major factors required for high oil accumulation of yellow nutsedge in relative to purple nutsedge.

### Transcripts for Carbon Metabolism Toward Fatty Acid Synthesis Were Slightly Higher in Yellow Nutsedge Than Purple Nutsedge

The generation of plastid pyruvate for FA synthesis from sucrose primarily involves sucrose degradation in cytosol, hexose glycolysis and pentose phosphate pathway (PPP) occurring in both cytosol and plastid (Fig. 3A, B).

**Fig. 3.**
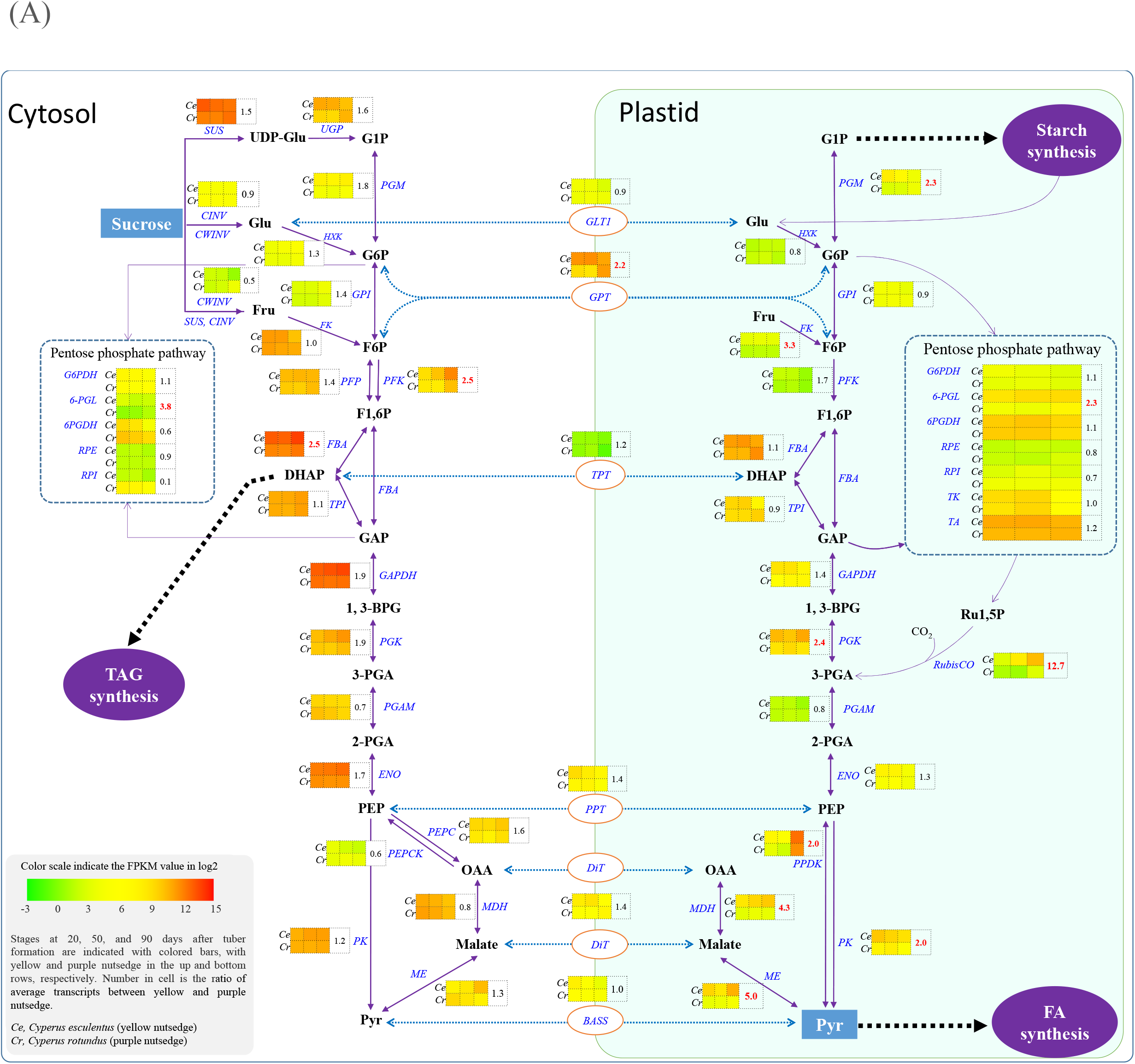
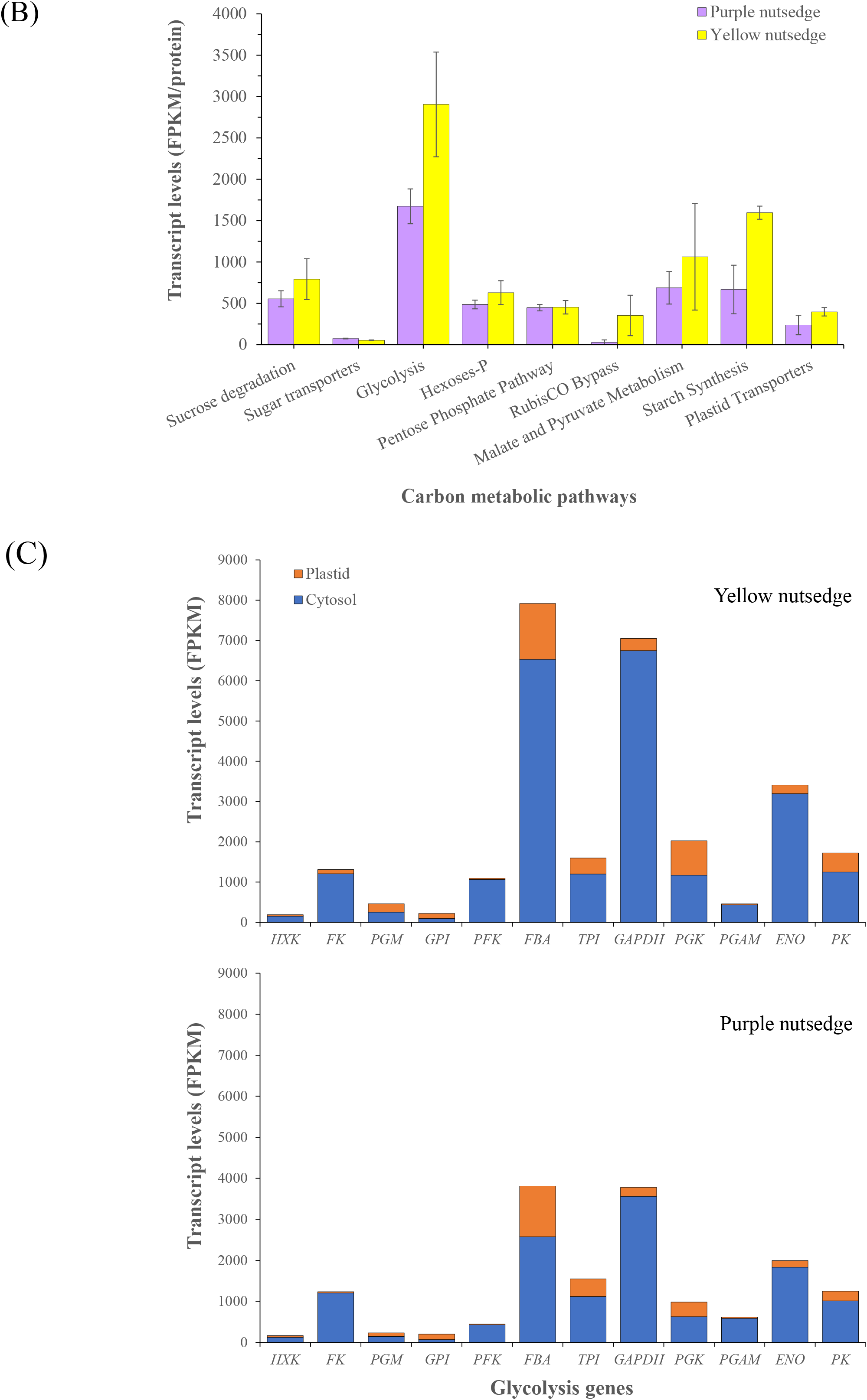
Transcript levels for carbon metabolism enzymes. (A) Transcript levels for various carbon metabolic pathways. The level represented by FPKM and the data are averaged on three developing stages of tubers, with error bars representing their standard deviation. Transcripts for proteins with isoforms or multiple subunits were summed. (B) Transcript patterns for genes involved in carbon metabolism. Gene names are indicated in blue. Value in table cell indicates the transcript ratio of yellow to purple nutsedge. Ratios without less than twofold are represented in red boldface. (C) Transcript levels for glycolysis enzymes in the cytosol and the plastid. Abbreviations: 1,3-BPG, 1,3-Bisphosphoglycerate; 2-PGA, 2-phosphoglycerate; 3-PGA, 3-phosphoglycerate; 6PGDH, 6-phosphogluconate dehydrogenase; 6-PGL, 6-phosphogluconolactonase; BASS, sodium bile acid symporter family protein; CINV, cytosolic invertase; CWINV, cell wall invertase; DHAP, dihydroxyacetone phosphate; ENO, enolase; F1,6P, fructose 1,6 bis-phosphate; F6P, fructose 6-phosphate; FBA, fructose bisphosphate aldolase; FK, fructokinase; Fru, fructose; G1P, glucose 1-phosphate; G6P, glucose 6-phosphate; G6PDH, glucose 6-phosphate dehydrogenase; GAP, glyceraldehyde 3-phosphate; GAPDH, glyceraldehyde 3-phosphate dehydrogenase; GLT1, plastidic glucose translocator 1; Glu, glucose; HXK, hexokinase; MDH, malate dehydrogenase; ME, malic enzyme; NTT, nucleoside triphosphate transporter; OAA, oxaloacetate; PEP, phosphoenol pyruvate.; PEPC, phosphoenolpyruvate carboxylase; PEPCK, phosphoenolpyruvate carboxykinase; PFK, phosphofructokinase; PFP, pyrophosphate dependent phosphofructokinase; PGAM, 2,3-bisphosphoglycerate-dependent phosphoglycerate mutase; PGI, phosphoglucose isomerase; PGK, phosphoglycerokinase; PGM, phosphoglucomutase; PK, pyruvate kinase; PPDK, pyruvate orthophosphate dikinase; PPT, phosphoenolpyruvate/phosphate antiport; PRK, phosphoribulokinase; Pyr, pyruvate; RCA, RubisCO activase; RPE, ribulose-phosphate 3-epimerase; RPI, ribose-5-phosphate isomerase; Ru1,5P, ribulose-1,5-bisphosphate; Rubisco, ribulose bisphosphate carboxylase; SUS, sucrose synthase; TA, transaldolase; TK, transketolase; TPI, triose phosphate isomerase; TPT, triosephosphate/phosphate antiport; UGP, UDP-glucose pyrophosphorylase.

It was found that there were only 1.4-fold higher transcripts in yellow nutsedge compared to purple nutsedge for sucrose degradation pathway (Fig. 3B), which is catalyzed either by sucrose synthase (SUS) into uridine diphosphate glucose (UDP-Glu) and fructose (Fru), or by extracellular cell wall invertase (CWINV) or intracellular neutral invertase (CINV) into glucose (Glu) and Fru (Fig. 3A). *SUS* genes in both nutsedges were highly expressed at levels of over 2800 FPKM/protein that were at least 15-fold higher than *CINV or CWINV* (Fig. 3A, Supplemental Table S2), implying that SUS might play an important role as the preferred enzyme in initial sucrose metabolism in nutsedge tubers. This result could support the evidence that SUS activities in plant seeds or potato tuber were significantly higher than INV activities (Xu et al, 1989; Appeldoorn et al, 1997; Ruuska et al, 2002; Morley-Smith et al, 2008).

Similar patterns of gene expresses involved in glycolysis were also present in the two tuber tissues (Fig. 3B). Transcripts for nearly all of glycolytic enzymes in cytosolic or plastidial compartments were at slightly higher or similar levels in yellow nutsedge compared with purple nutsedge (Fig. 3A). Only expression levels for cytosol isoforms of ATP-dependent phosphofructokinase (PFK) and fructose-bisphosphate aldolase (FBA), and plastid fructose kinase (FK), which were over 2.5-fold higher in yellow nutsedge than in purple nutsedge.

Comparing the transcripts for both plastid and cytosol glycolysis revealed some conserved features between the two tuber tissues. The transcript levels were much higher for almost all genes involved in cytosolic glycolysis than their counterparts for plastidial glycolysis (Fig. 3C), except for genes encoding for phosphoglucomutase (PGM) and glucose-6-phosphate isomerase (GPI) where transcripts were distributed in somewhat balanced manner between the two glycolytic pathways. For a complete glycolytic pathway in the two plant tubers, the glycolytic genes involved in late glycolysis associated with the seven downstream glycolytic steps (from fructose-bisphosphate aldolase (FBA) to pyruvate kinase (PK)) were overall more abundantly expressed in relative to those in early glycolysis. Notably among these genes, those encoding for cytoplasmic glyceraldehyde-3-phosphate dehydrogenase (GAPDH), FBA and enolase (ENO) were significantly highly expressed (>1,800 FPKM). As a result, these data suggest that the cytosolic glycolysis metabolic pathway, particularly the late process, is highly active and might produce most of carbon precursors toward FA synthesis in the two tubers. The gene expression patterns of these two species resemble those observed in heterotrophic non-green oil seeds such as castor, safflower and sunflower (Troncoso-Ponce et al, 2011), but differ from those in photoheterotrophic green seeds such as Arabidopsis, rapeseed and soybean (Agrawal et al, 2008) and oil-rich mesocarps of oil palm as well avocado that displayed more balanced distribution of transcripts between cytosol and plastid glycolysis (Bourgis et al, 2011; Dussert et al, 2013; Kilaru et al, 2015).

Pentose phosphate pathway, a glycolysis bypass process, has been shown to provide carbon sources for pyruvate generation, which also occurred in both cytosol and plastid (Baud and Lepiniec, 2010). However, the overall transcripts for this pathway in cytosol or plastid is not much different between the two species (Fig. 3B). Only a significant increase in cytosolic ribose-5-phosphate isomerase (RPI) enzyme was observed in purple nutsedge compared with yellow nutsedge (Fig. 3A).

### Significant Difference of Transcripts for Plastid Rubisco Bypass Is Present Between Two Species

Glycolysis is also bypassed by the production of 3-phosphoglycerate (3-PGA) catalyzed by ribulose-1,5-bisphosphate carboxylase/oxygenase (RubisCO), a key enzyme for fixing carbon dioxide. This process without full Calvin cycle is called Rubisco bypass or shunt (Schwender et al, 2004). Intriguingly, the expression of gene encoding for RubisCO ortholog was also appeared in two nutsedge tubers. In this study, six unigenes encoding for RubisCO small subunit (RbcS) and one for RubisCO large subunit (RbcL, ATCG00490) were detectable to express during tuber development, though RbcL was merely slightly transcribed (Fig. 4A; Supplemental Table S2). The presence of RubisCO genes, particularly *RbcS* in nutsedge tubers is surprising, since tubers are non-green and non-photosynthetic underground tissues that require little or no light for their development and growth. It was also found that RbcS homologous were expressed exclusively in other non-photosynthetic tissues including seeds, fruits, roots, and secretory organs such as trichomes, but seem to be almost absent in photosynthetic tissues (Morita et al, 2014; Morita et al, 2016). It is unclear why these RbcS proteins are maintained in the non-photosynthetic tissues including tuber, but one can expect that this type of RbcS may be evolved to accomplish different functions other than Calvin cycle occurred in photosynthetic tissues. It suggested that Rubisco, particularly RbcS, plays a specialized role in recycling of CO_2_ at high concentration environment generated by the metabolism in tissues that are less permeable to gas exchange (Pottier et al, 2018).

**Fig. 4.**
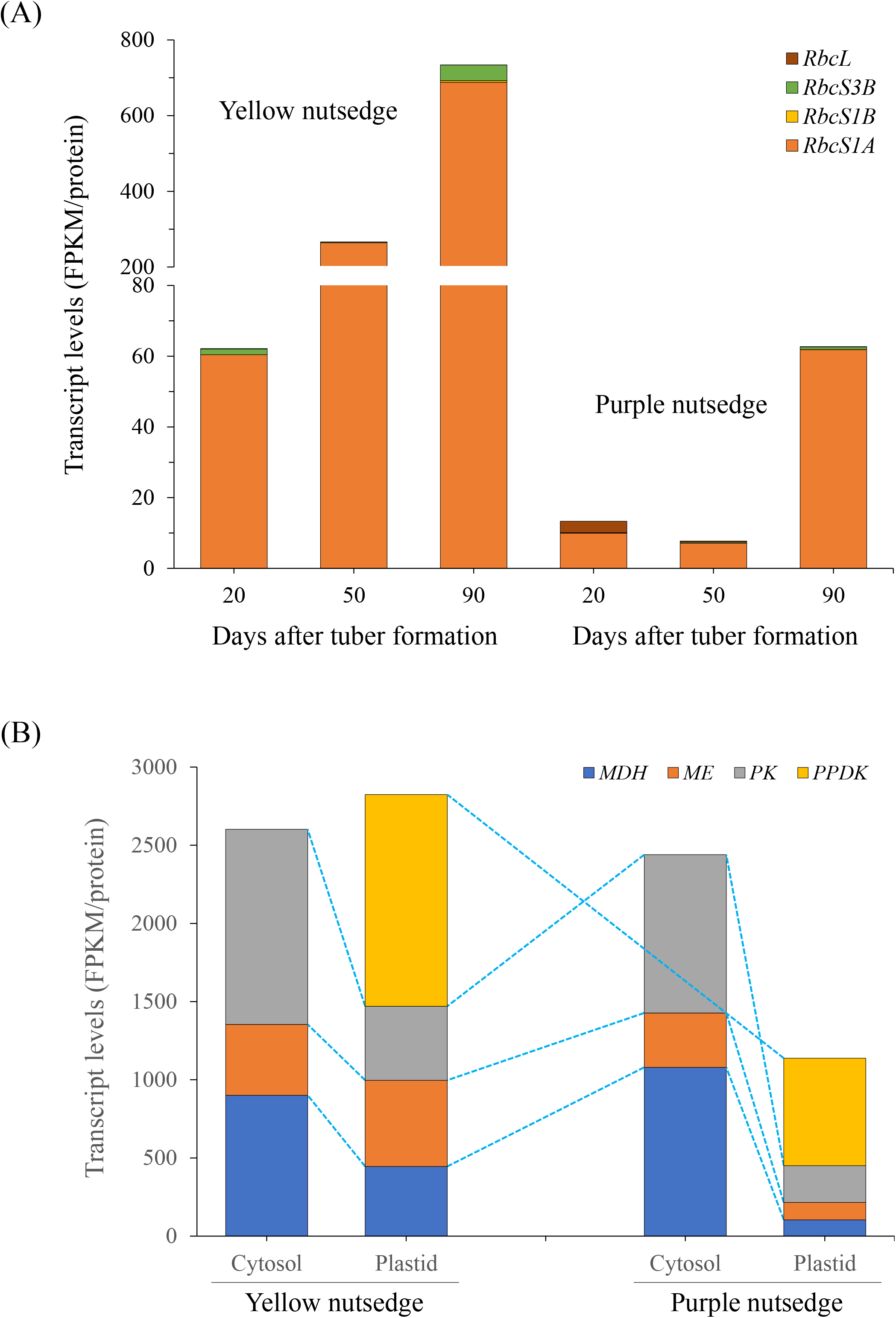
Transcript levels for RubisCO bypass (A) and pyruvate generation. (A) Temporal changes in transcript levels for RubisCO small and large subunits. (B) Transcript levels for malate and pyruvate metabolism enzymes.

Comparatively, the RbcS genes in purple nutsedge is poorly transcribed at low levels, while they were abundantly expressed in yellow nutsedge and displayed upregulation patterns during tuber development (Fig. 4A), with the average transcript levels being more 12 times higher than in purple nutsedge (Fig. 3A). A recent study also showed that the ortholog of RbcS in potato starchy tuber displayed much more abundant transcripts in oil-accumulated transgenic lines expressing Arabidopsis *WRI1* gene than in the control (Hofvander et al, 2016). These results might reinforce the enzyme assay of the silique of rapeseed, which revealed that high oil content was associated closely with enhanced *RbcS* expression levels and its activities (Hua et al, 2012). Evidence has shown that changes in the RbcS transcript abundance were directly correlated with the changes in Rubisco level (Izumi et al, 2010; Atkinson et al, 2017). High expression of RbcS genes possibly reflected the high CO_2_ environment (Ruuska et al, 2004) and was most likely associated with the ability of capturing CO_2_ resulting from the conversion of malate to pyruvate by the plastid NADP-dependent malic enzyme (ME) or pyruvate to acetyl-CoA by the plastid pyruvate dehydrogenase complex (PDHC) (Ruuska et al, 2002). Previous studies demonstrated that RubisCO bypass or shunt skipping Calvin cycle for CO_2_ recapture in green oil-rich seeds brought about less loss of carbon as CO_2_ and produced more 3-PGA toward for FA synthesis, thus improving carbon conversion efficiency (Ruuska et al, 2004; Schwender et al, 2004; Alonso, et al, 2007; Allen et al, 2009; Tsogtbaatar et al, 2020).

Whether the expression of *RbcS* gene is also related to high CO_2_ concentration and involved in recycling CO_2_ in tubers remains to be further studied. Nevertheless, the relatively high expression level of *RbcS* in yellow nutsedge compared to purple nutsedge indicated that RbcS might play a part role in carbon metabolism for oil production in oil tuber tissue.

### Transcripts for Plastid Malate and Pyruvate Metabolism Are Strikingly Distinct Between Two Species

Pyruvate is the important carbon precursor required for plastid fatty acid synthesis. It can be generated either from phosphoenolpyruvate (PEP) catalyzed by PK or pyruvate phosphate dikinase (PPDK), or through the malate metabolism involved in malate dehydrogenase (MDH) and ME that catalyze the sequential production of malate and pyruvate from oxaloacetate (OAA) (Fig. 3A). In this study, the abundant expression of genes encoding for MDH, ME, and PK occurred in both cytosol and plastid, and transcript levels for MDH and ME were comparable to those for PK (Fig. 4B), implying that the malate metabolism is as important as PK in pyruvate generation and pyruvate is produced from both hexose via the glycolytic pathway and NADP-dependent malic enzyme for fatty acid synthesis in nutsedge tubers. Previous studies showed that malate was a major substrate for fatty acid synthesis in non-green seeds of safflower, castor bean, and sesame (Browse and Slack, 1985; Eastmond et al, 1997; Suh et al, 2003)

Higher expression of MDH, ME, and PK was noted in the cytosol than in the plastid (Fig. 4B), suggesting that the cytosolic pyruvate generation might be more prominent in two species, and malate and pyruvate produced in the cytosol could be transported to the plastid for subsequent pyruvate metabolism for fatty acid synthesis.

One interesting aspect from this comparison was that transcripts for cytosolic MDH, ME and PK were comparable in both nutsedge species (Fig. 4B). By contrast, transcript levels for plastidic counterpart and PPDK enzymes were more than two times higher in yellow nutsedge than in purple nutsedge. This suggests that plastid malate and pyruvate metabolism was more active and might produce more carbon source as well as reducing power (NADH/NADPH) required for fatty acid synthesis in yellow nutsedge relative to purple nutsedge. Therefore, our results imply that up-regulation of genes involved in plastid malate and pyruvate metabolism plays an important role in providing pyruvate for high oil synthesis.

### Transcripts for Fatty Acid Synthesis Enzymes in Plastid Were More Abundant in Yellow Nutsedge than in Purple Nutsedge

The expressions of at least fourteen proteins required for *de novo* fatty acid synthesis in the plastid from pyruvate were all detectable in two species (Fig. 5A). Among these enzymes/proteins, PDHC, acetyl-CoA carboxylase (ACCase), and acyl-carrier protein (ACP) transcribed more abundant than any other enzymes (Fig. 5B), implicating the important roles of these proteins in fatty acid synthesis. The overall transcript levels for these three proteins accounted for over 50% of total for FA synthesis at each stage of tuber development. Similar phenomena were also observed in developing oil-rich seeds and mesocarps (Troncoso-Ponce et al, 2011; Bourgis et al, 2011; Kilaru et al, 2015).

**Fig. 5.**
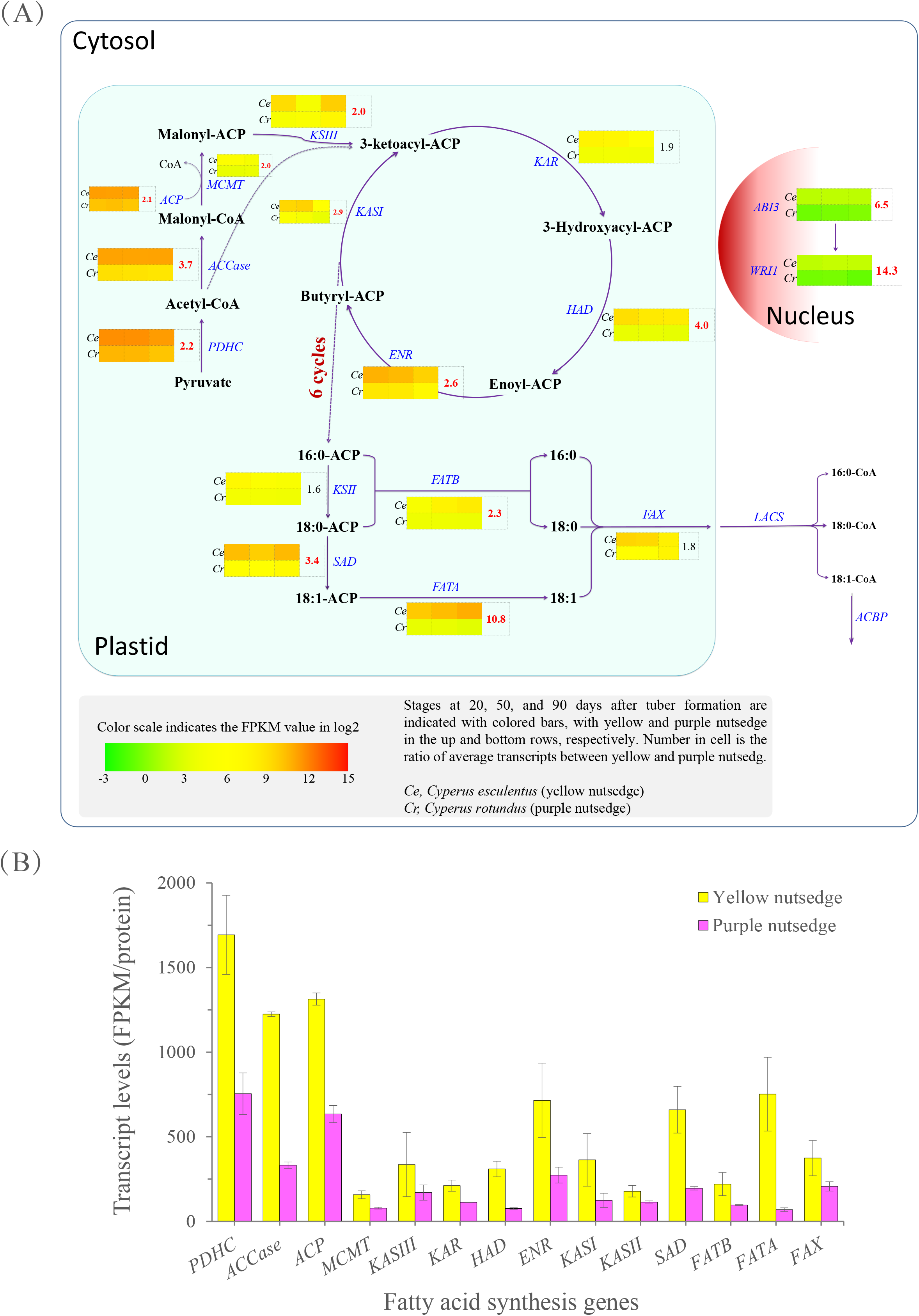
Transcript levels of fatty acid synthesis. (A) Schematic pathway of fatty acid synthesis. Enzyme or protein names are indicated in blue. Gene names are indicated in blue. Value in table cell indicates the transcript ratio of yellow to purple nutsedge. Ratios more than twofold are showed in red boldface. (B) Average transcript levels for plastid fatty acid synthesis genes. The data are averaged on three tuber developing stages, with error bars indicating standard deviation. Abbreviations: ABI3, Abscisic Acid Insensitive 3; ACBP, acyl-CoA-binding protein; ACP, Acyl Carrier Protein; ACCase, acetyl-CoA carboxylase; ENR, Enoyl-ACP Reductase; FAT, Acyl-ACP Thioesterase; FAX, Fatty Acid Export; HAD, Hydroxyacyl-ACP Dehydratase; KAR, Ketoacyl-ACP Reductase; KAS, Ketoacyl-ACP Synthase; LACS, Long-Chain Acyl-CoA Synthetase; MCMT, Malonyl-CoA:ACP Malonyltransferase; PDHC, Pyruvate Dehydrogenase Complex; SAD, Stearoyl-ACP desaturase; WRI1, WRINKLED1

Almost all proteins were transcribed at comparatively higher levels in yellow nutsedge, coinciding with high oil accumulation in its tubers. Overall, transcript levels for these plastidial proteins were on average 2.5-fold higher in yellow nutsedge than in purple nutsedge (Fig. 2C). Significant individual differences were present for ACCase, hydroxyacyl-ACP dehydratase (HAD), stearoyl-ACP desaturases (SAD), and acyl-ACP thioesterase A (FATA), for which the transcripts were more than 3-fold higher in yellow nutsedge compared with purple nutsedge (Fig. 5A), suggesting that the metabolic pathways catalyzed by these enzymes perhaps were much more active in yellow nutsedge.

ACCase in the plastid is one of the main enzymes in regulating oil production, as it catalyzes the first truly committed step of the irreversible carboxylation of acetyl-CoA to malonyl-CoA with no other known metabolic fate, a rate-limiting pathway in the control of fatty acid synthesis (Sasaki and Nagano, 2004). The importance of ACCase in the control of plant lipid synthesis was confirmed by the evidence that about 55% of the total carbon flux for overall fatty acid synthesis was controlled by the ACCase in leaf tissues (Page et al, 1994). In this study, ACCase transcripts represented about 15% and 11% of the total levels for fatty acid synthesis enzymes in yellow nutsedge and purple nutsedge, respectively. Among the four subunits of heteromeric ACCase enzyme complex, three nuclear-coded subunits, biotin carboxylase (BC), followed by biotin carboxy-carrier protein (BCCP) with α-carboxyltransferase (α-CT), were most abundant in two species, while the plastid-coded subunit (β-carboxyltransferase, β-CT) was barely transcribed at very low levels (<3 FPKM) (Fig. 6A). The relatively higher transcripts for nuclear-coded subunits against plastid-coded ones were also observed in oil-rich seeds and fruits (Troncoso-Ponce et al, 2011; Bourgis et al, 2011; Kilaru et al, 2015), suggesting a conserved major role of nuclear-coded subunits for ACCase activity in diverse plants and tissues. In case of nuclear-coded three ACCase, transcripts for each subunit were 3.2 to 4.5 times higher in yellow nutsedge than in purple nutsedge. Previous study indicated that high ACCase mRNAs were consistent with actively synthesizing fatty acids and accumulating high levels of oil in seed (Ke et al, 2000). Therefore, our results implied that the initial reaction of fatty acid synthesis catalyzed by ACCase, with the three ACCase subunits in a coordinated expression pattern, was more active and might provide more malonyl-CoA substrate needed for oil production in yellow nutsedge as to purple nutsedge.

**Fig. 6.**
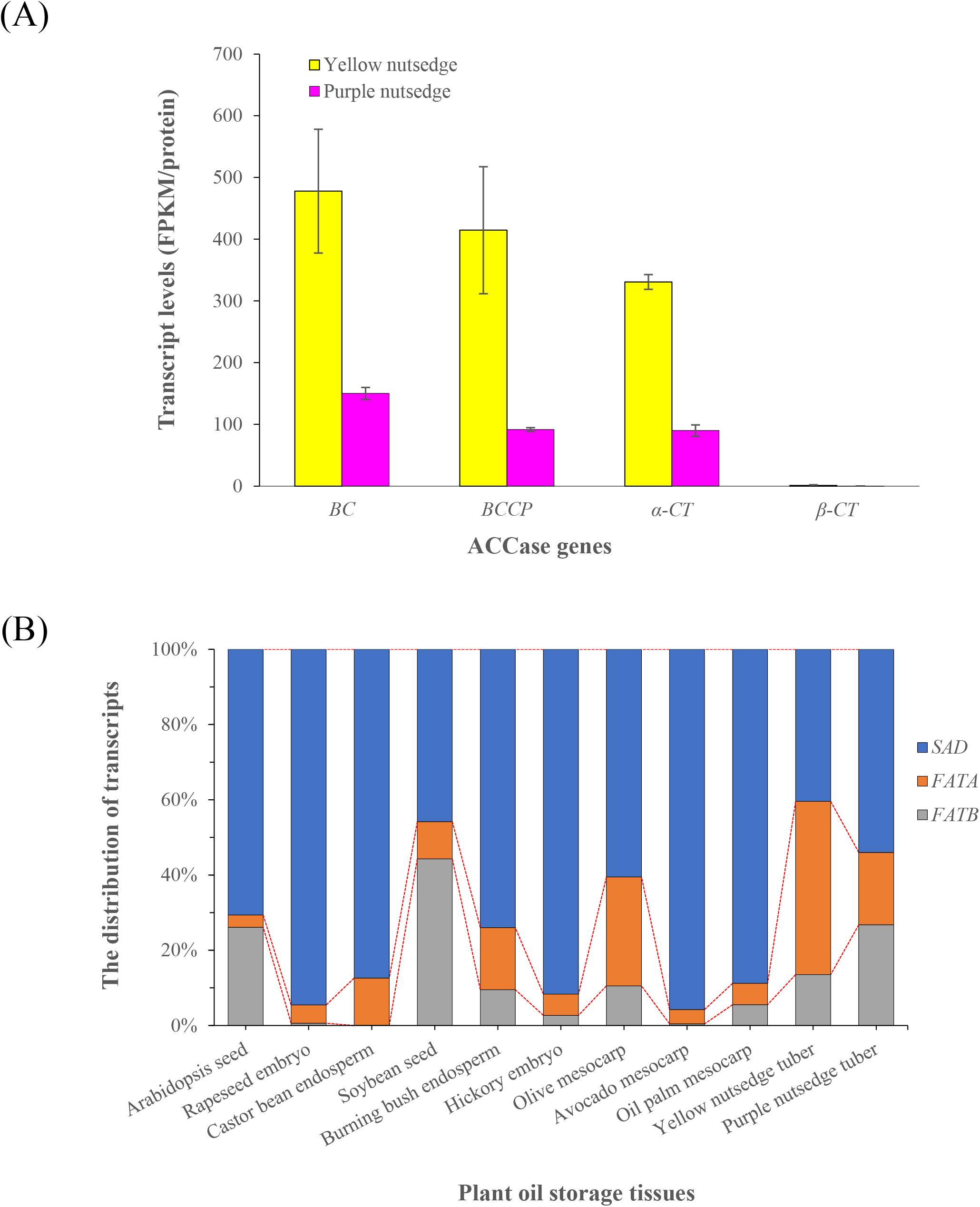
Expression pattern for ACCase (A) and acyl-ACP desaturase and thioesterase (B). (A) Average transcript levels for diverse ACCase genes. The data are averaged on three tuber developing stages, with error bars indicating standard deviation. (B) Relative distribution of transcript levels for SAD, FATA and FATB in plant oil-rich tissues. The data are averaged on all the developing stages of sink tissues. The transcript values for subunits of a protein and for multiple isoforms were summed.

HAD is responsible for the dehydration of 3-hydroxyacyl-ACP to generate trans-2-enoyl-ACP. In Arabidopsis, two genes (At2g22230 and At5g10160) have been identified to code for HAD, and they were highly expressed during fatty acid synthesis in developing seeds (Schmid et al, 2005). While the transcripts for the two *HAD* genes were detectable in most oil-rich plants (Bourgis et al, 2011; Troncoso-Ponce et al, 2011; Jones and Vodkin, 2013; Bertioli et al, 2016; González-Thuillier et al, 2016), only one *HAD* ortholog (At5g10160) was represented in the two nutsedge tubers, similar to the case observed in other underground sink tissues such as potato (Hofvander et al, 2016), sweet potato (Tao et al, 2017), cassava (Bredeson et al, 2016) and sugar beet (Bellin et al, 2005), suggesting its crucial role in fatty acid synthesis and a housekeeping function at least in underground storage tissues. A single HAD ortholog was also noted for other oil-rich plants, for example, castor bean and olive (*Olea europaea*) contain one At5g10160 ortholog (Troncoso-Ponce et al, 2011; Unver et al, 2017), while avocado include only At2g22230 ortholog (Kilaru et al, 2015). These results indicated that the expressions of HAD orthologs are species- and tissue-specific, and different orthologs may have evolved to respond to fatty acid synthesis in different plants.

SAD and FATA are two important enzymes controlling levels of unsaturated fatty acid. High expression of *SAD* along with *FATA* in relative to *FATB* was found to occur in unsaturated fatty acid-rich oil tissues including seeds, fruits and tubers (Fig. 6B). During the tuber development, transcripts for *SAD* or *FATA* were much abundant in yellow nutsedge relative to purple nutsedge; particularly, *FATA* genes were up-regulated during development and transcribed at more than 10-fold higher levels in yellow nutsedge than in purple nutsedge at tuber maturation, associating with their fatty acid profiles of oil. It is noteworthy that in yellow nutsedge, transcripts for FATA were 3-fold more abundant as compared to FATB. In contrast, the FATA transcript levels were less than and comparable to FATB in purple nutsedge. Intriguingly, high ratio of FATA to FATB transcript was also observed in other plant oil storage tissues rich in oleic acid or its derivatives, such as oil seeds of rapeseed and castor bean (Troncoso-Ponce et al, 2011) as well as hickory (Huang et al, 2016), and oil fruits of avocado (Kilaru et al, 2015) and olive (Alagna et al, 2009; Unver et al, 2017). By contrast, the transcript ratio of FATA/FATB is only 0.1, 0.2, and 1.0, respectively, for seeds of Arabidopsis (Troncoso-Ponce et al, 2011) and soybean (Jones and Vodkin, 2013) and mesocarp of oil palm (Bourgis et al, 2011) that contain low levels of oleic acid. Altogether, these results revealed that the expression patterns of SAD and FATA genes in two tubers reflect the fatty acid composition, and the ratio of FATA to FATB expression is most likely correlated with the content of C18:1.

### Transcripts for Most TAG Synthesis Genes in Endoplasmic Reticulum Were Similar or Less in Yellow Nutsedge Than in Purple Nutsedge

In contrast to transcript patterns for plastidial fatty acid genes, most TAG synthesis genes in endoplasmic reticulum were expressed at similar or less levels in yellow nutsedge as compared to purple nutsedge (Fig. 7A). For example, phosphatidic acid phosphohydrolase (PAP), phospholipid:diacylglycerol acyltransferase (PDAT), lysophosphatidylcholine acyltransferase (LPCAT) and Δ12-oleate desaturase (FAD2), an enzyme responsible for synthesis of C18:2 fatty acid in the endoplasmic reticulum, showed similar expression patterns between yellow and purple nutsedge, while lysophosphatidyl acyltransferase (LPAAT) and two enzymes that catalyze the exchanges of fatty acyl groups between phosphocholine (PC) and diacylglycerol (DAG), cytidine-5-diphosphocholine: diacylglycerol cholinephosphotransferase (CPT) and phosphatidylcholine: diacylglycerol cholinephosphotransferase (PDCT), had 2- to 5-fold lower transcripts in yellow nutsedge than in purple nutsedge.

**Fig. 7.**
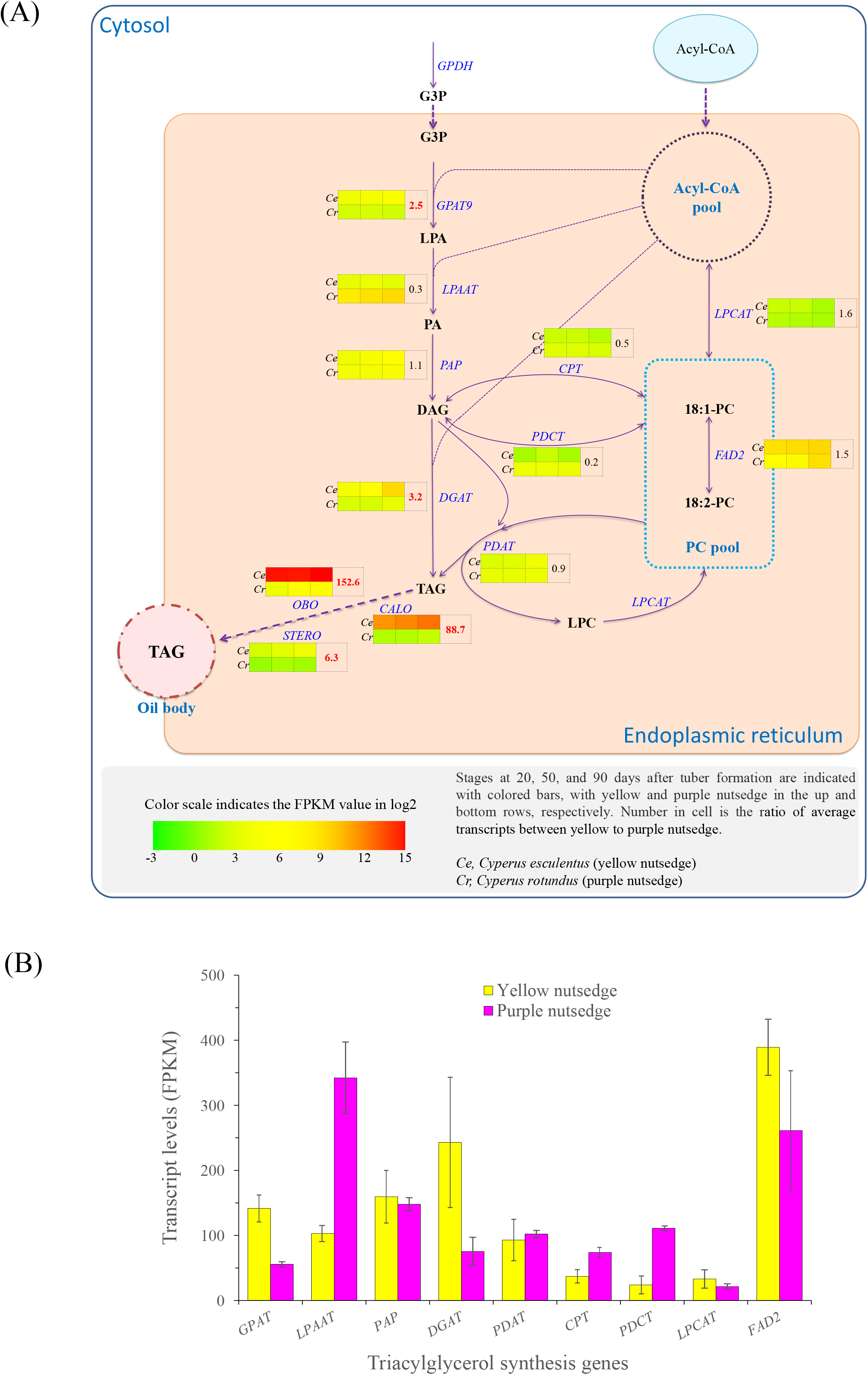
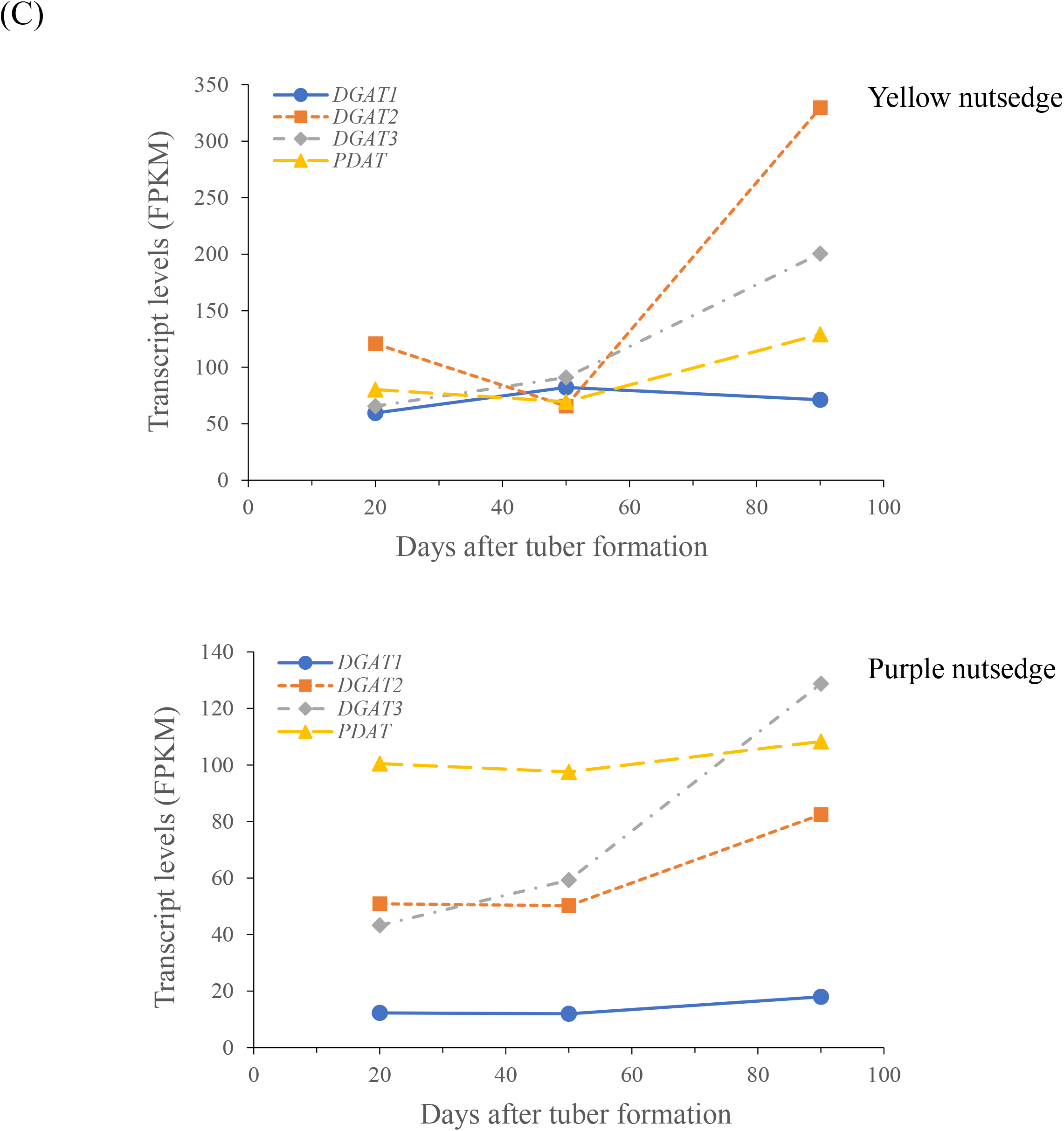
Expression patterns for genes related to triacylglycerol synthesis. (A) Schematic of triacylglycerol (TAG) synthesis pathways. Gene names are indicated in blue color. (B) Average transcript levels for TAG synthesis genes in two nutsedges. The data are averaged on all three developing stages of tubers. The transcript values for subunits of a protein and for multiple isoforms were summed. (C) Temporal transcripts for DGAT and PDAT enzymes during tuber development. Abbreviations: CPT, Diacylglycerol cholinephosphotransferase; DAG, Diacylglyceol; DGAT, Acyl-CoA:Diacylglycerol Acyltransferase; FAD2, Oleate desaturase, Fatty acid desaturase 2; G3P, Glycerol-3-Phosphate; GPAT, Glycerol-3-Phosphate Acyltransferase; GPDH, NAD-dependent Glycerol-3-Phosphate Dehydrogenase; LPA, lysophosphatidic Acid; LPAAT, 1-acylglycerol-3-phosphate acyltransferase; LPC, lysophosphatidylcholine; LPCAT, 1-acylglycerol-3-phosphocholine Acyltransferase; PA, Phosphatidic Acid; PAP, Phosphatidic Acid Phosphohydrolase; PC, Phosphatidylcholine; PDAT, Phospholipid:Diacylglycerol Acyltransferase; PDCT, Phosphatidylcholine:diacylglycerol cholinephosphotransferase;

Two exceptions were noted for glycerol-3-phosphate acyltransferase (GPAT9) and diacylglycerol acyltransferase (DGAT) that presented 2.5- and 3.4-fold higher transcripts in yellow nutsedge than in purple nutsedge, respectively (Fig. 7). DGAT is the key enzyme that catalyzes the final step of acyl-CoA dependent TAG synthesis (Liu et al. 2012). In the two species, transcripts were much higher for DGAT2 than for DGAT1, suggesting a more prominent role of DGAT2 than DGAT1, and DGAT2 may be a key mediator in tuber oil production. Our recent study of DGAT1 and DGAT2 functional analysis has provided an evidence to support this hypothesis (Liu et al, 2020). It was noted that PDAT, an enzyme responsible for the transfer of fatty acyl moiety from PC to DAG destined to TAG synthesis, and an ortholog of *Arabidopsis thaliana* DGAT3 (At1g48300), the soluble cytosolic enzyme that might catalyze TAG synthesis using cytosolic acyl-CoA pool (Hernández et al, 2012), were also detectable to have more abundant transcripts than DGAT1 during tuber maturation (Fig. 7C), but displayed similar expression patterns between the two species. Furthermore, transcripts for DGAT2, DGAT3 and PDAT enzymes in yellow nutsedge were all increased during tuber maturation, coinciding with tuber oil accumulation. Taken together, our data implicate the important roles of DGAT2, DGAT3, and PDAT rather than DGAT1 played in transcriptional regulation of TAG synthesis in the nutsedge tubers.

### Great Transcriptional Divergence of TAG Storage Genes Between Two Species

Similar to oil seeds of plants (Troncoso-Ponce et al, 2011), oil-tuber of yellow nutsedge were represented by the abundant expression of a large number of genes encoding for seed-like oil body proteins such as oleosin (OBO), caleosin (CALO), steroleosin (STERO), oil body associated protein (OBAP), and seed lipid droplet protein (SLDP), with the OBO transcript being the most abundant (Fig. 8; Supplemental Table S3). In addition, their expression levels were increased during tuber maturation, consistent with the tuber oil accumulation. Large difference was noted for these proteins, for which their transcripts were over 6- to 160-fold higher in yellow nutsedge than in purple nutsedge. Similar contrast was also detectable across all the developmental stages. Therefore, all these results might imply the important roles of these oil body proteins particularly oleosin in stabilizing TAG and producing high oil content in yellow nutsedge. Previous studies have shown that oleosin is accumulated in a coincident manner with TAG accumulation and the abundant expression of oleosin is associated with high oil content in seeds (Frandsena et al, 2001; Miquel et al, 2014; Zhang et al, 2016; Huang, 2018; Xiao et al, 2019; Ischebeck et al, 2020).

**Fig. 8.**
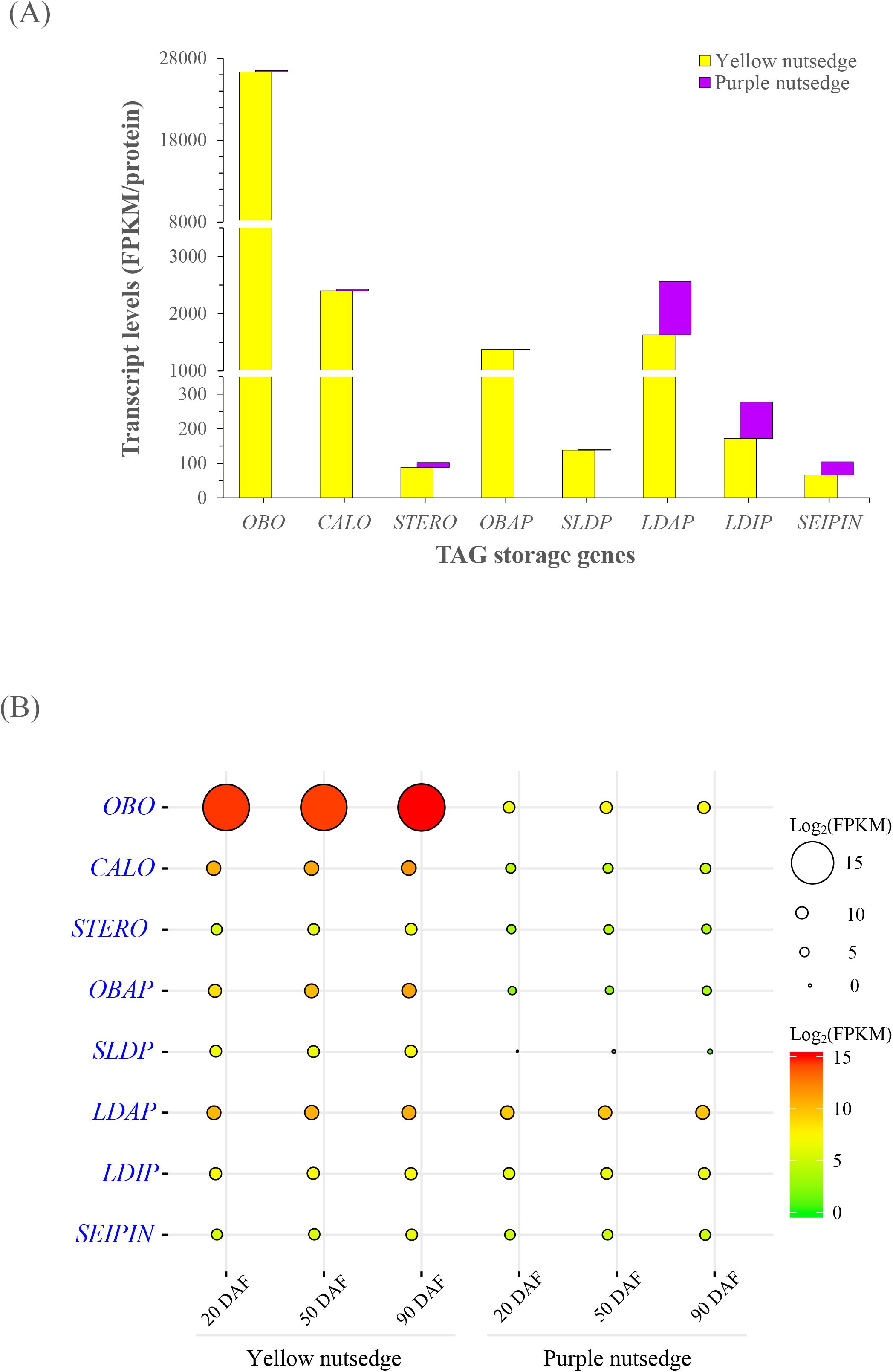
Transcript patterns for triacylglycerol storage proteins. (A) Average transcript levels for diverse TAG storage genes. The data are averaged on three tuber developing stages and the transcript values for subunits of a protein and for multiple isoforms were summed. (B) Balloon plot showing temporal changes in transcript levels (as log2(FPKM)) for TAG storage genes. Abbreviations: CALO, Caleosin; LDAP, Lipid droplet associated protein; LDIP, LDAP-interacting protein; OBAP, Oil body-associated protein; OBO, Oleosin; SEIPIN, adipose-regulatory protein.; SLDP, Seed lipid droplet protein; STERO, Steroleosin. DAF, days after tuber formation.

The high transcripts of oil body proteins in yellow nutsedge tuber are in sharp contrast to non-seed oily mesocarp tissues of olive, oil palm and avocado that showed lowly expressed transcripts for these structural proteins (Ross et al, 1993; Giannoulia et al, 2007; Bourgis et al, 2011), which were considered less contribution to TAG storage or assembly in oil mesocarp tissues. Indeed, other lipid droplet protein such as lipid droplet-associated protein (LDAP) were found to display plentiful transcripts in these oil mesocarps (Bourgis et al, 2011; Kilaru et al, 2015).

Intriguingly, a number of genes encoding for LDAP (At3g05500), LDAP-interacting protein (LDIP, At5g16550) and lipodystrophy protein (SEIPIN1, AT5G16460; SEIPIN2, AT1G29760) were also detectable to moderately express in developing tubers of two species (Fig. 8; Supplemental Table S3). The transcript level for LDAP or LDIP were even higher than that of STERO. In addition, the temporal transcript patterns of the three proteins were similar to those of seed-like oil body proteins and were in accordance with oil accumulation during tuber development. However, transcripts for the three proteins showed no substantial differences between yellow and purple nutsedge.

It is noteworthy that among these eight proteins, OBO was transcribed most abundantly in yellow nutsedge, followed by CALO and LDAP in this descending order, similar to the case in oil seeds where they are predominantly expressed and the most abundant proteins on oil body (Huang, 2018). By contrast, LDAP showed highest transcript level in purple nutsedge, which are 5 to 66 times higher than those of other oil body proteins.

Taken together, our results described above indicated that expression patterns of oil body proteins involved in TAG storage is positively associated with oil accumulation, but transcriptional control of these proteins is significantly different between the two species, most possibly tissue-specific.

### WRI1 and ABI3 Show Much Higher Transcripts in Yellow Nutsedge Than in Purple Nutsedge

WRINKLED1 (WRI1) is an important transcriptional regulator in controlling oil accumulation in aboveground oil-rich seeds and fruits of plants, and has been well functionally characterized particularly in the model plant *Arabidopsis thaliana* (Cernac and Benning 2004; Baud et al. 2009). In the two nutsedge species, *WRI1* and its multiple orthologs, *WRI2* (AT2G41710), *WRI3* (AT1G16060), and *WRI4* (AT1G79700), belonging to the APETALA2-ethylene responsive element binding protein (AP2/EREBP) family, were all detectable to express in tubers, but they displayed low transcript levels (<25 FPKM) (Fig. 9). Particularly, *WRI3* and *WRI4* were only barely expressed, suggesting that they are unlikely to participate in the control of oil production in tubers. This may support the fact that in Arabidopsis, only *WRI1* can activate fatty acid synthesis in oil seeds for oil production, and *WRI3* and *WRI4* are required for cutin synthesis in floral and stem tissues (To et al, 2012). Although *WRI2* transcript levels were relatively higher compared to *WRI3* and *WRI4*, they were comparable between yellow and purple nutsedge. *WRI2* was reported to was unlikely associated with fatty acid synthesis (To et al, 2012).

**Fig. 9.**
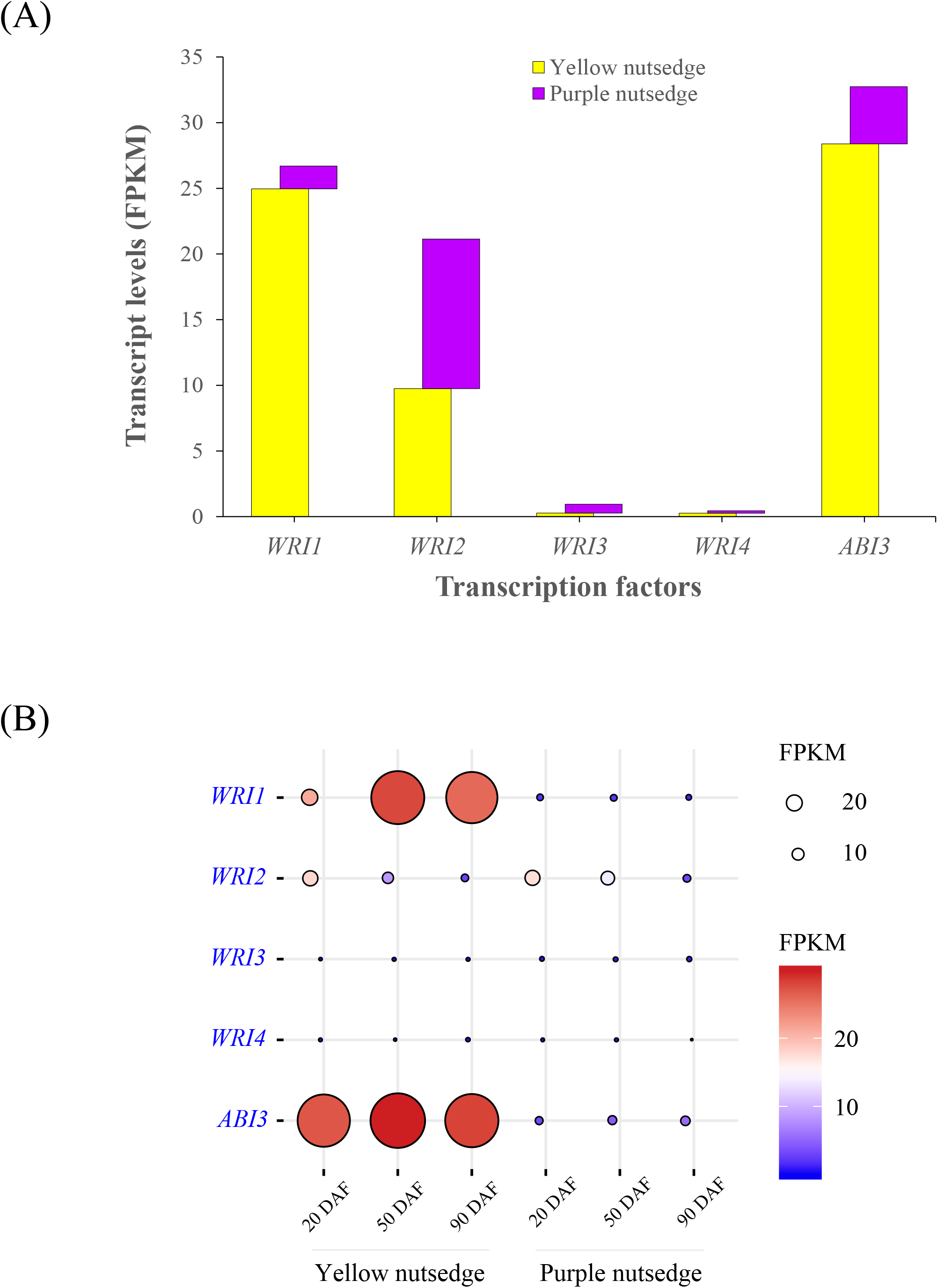
Transcript patterns of WRI1-like and ABI3 transcription factors. (A) Average transcript levels for WRI1-like proteins and ABI3 in nutsedge tubers. The data are averaged on three tuber developing stages and the transcript values for subunits of a protein and for multiple isoforms were summed. (B) Temporal changes in transcript levels (FPKM) for five transcription factors.

A significant difference between two species was noted for *WRI1* ortholog that showed 14.3-fold on average higher transcripts in yellow nutsedge over purple nutsedge (Fig. 9; Supplemental Table S3). A similar great contradiction was also observed between oil palm and date palm, in which oil palm mesocarp showed 57-fold higher expression of *WRI1* orthologue (Bourgis et al, 2011). In yellow nutsedge, the expression level of *WRI1* was increased with tuber development, which is in accordance with oil accumulation pattern in tubers (Fig. 9). Furthermore, the temporal expression pattern of *WRI1* matched that of its potential targets such as *PDH-β* (AT2G34590), *BCCP1* (AT5G16390), *FAB2* (AT2G43710), *FATA* (AT3G25110) and *FATB* (AT1G08510).

Unlike oil seed tissues, the maturation master regulators that directly or indirectly control the expression of *WRI1*, such as LEAFY COTYLEDON (LEC1 (AT1G21970) and LEC2 (AT1G28300)) and FUSCA3 (FUS3, AT3G26790) (Baud and Lepiniec, 2010), were not detectable in two nutsedge tubers, as in oil mesocarps of oil palm and avocado (Bourgis et al, 2011; Kilaru et al, 2015), supporting the previous report that FUS3-type proteins are present only in seed plants while LEC1- and LEC2-like proteins are appeared only in vascular and dicot plants, respectively (Li et al, 2010). Lack of these seed-like regulators suggested that the regulation of WRI1-related transcriptional network in non-seed tissues is different from that of oil seed tissues.

It was noteworthy that in yellow nutsedge tuber, an ortholog of *ABSCISIC ACID INSENSITIVE* 3 (*ABI3*) (AT3G24650), the member of B3 domain superfamily, showed transcript levels comparable to *WRI1* and quite similar temporal expression pattern (Fig. 9). Similar to WRI1, ABI3 ortholog was up-regulated significantly in yellow nutsedge as compared to purple nutsedge. Genes that share similar expression patterns are likely to interact with each other and have regulatory relationships of functionally importance (Ge et al, 2001). In this respective, however, it is unclear whether ABI3 is the upstream regulator of WRI1 and controls *WRI1* expression in the nutsedge tubers as in plant oilseeds. Recent study indicated that ABI3 played an important role in regulating plant oil accumulation, which might be independent from WRI1 and LEC2 regulation networks occurred in oil-rich seed tissues (Pouvreau et al, 2020).

To determine the relevant relations between *WRI1* and *ABI3* in regulating oil accumulation in oil tuber, a gene co-expression network was constructed and analyzed using the method of weighted gene co-expression network analysis (WGCNA) (Zhang and Horvath, 2005; Langfelder and Horvath, 2008). The produced network indicated that these two transcriptional factors were classified as hub genes within a same cluster or module in the co-expression network (data not shown). An interesting finding from the co-expression analysis is that *ABI3* and *WRI1* are co-expressed in concert with the oil genes involved in carbon metabolism, fatty acid synthesis, TAG synthesis, and TAG storage pathways (Fig. 10; Supplemental Table S4). Our analysis result of *WRI1* in concerted regulation with the genes related to carbon metabolism, fatty acid synthesis, and TAG synthesis in yellow nutsedge tuber is similar to recent reports (Guerin et al, 2016; Kuczynski et al, 2020). These oil-related target genes are co-regulated by both of *ABI3* and *WRI1*, indicating an overlapping set of targets for the two transcription factors. It was also shown that transcriptional regulation of *ABI3* and *WRI1* is closely interconnected and *WRI1* is one of direct targets for *ABI3*, suggesting that ABI3 is the regulator of WRI1 in yellow nutsedge, in contrast to oil-rich mesocarps of oil palm and avocado where WRI1 is not under the control of ABI3 (Bourgis et al, 2011; Kilaru et al, 2015). In oil palm, WRI1 transcriptional activation can be activated by three ABA-responsive transcription factors, NF-YA3, NF-YC2 and ABI5 (Yeap et al, 2017).

**Fig. 10.**
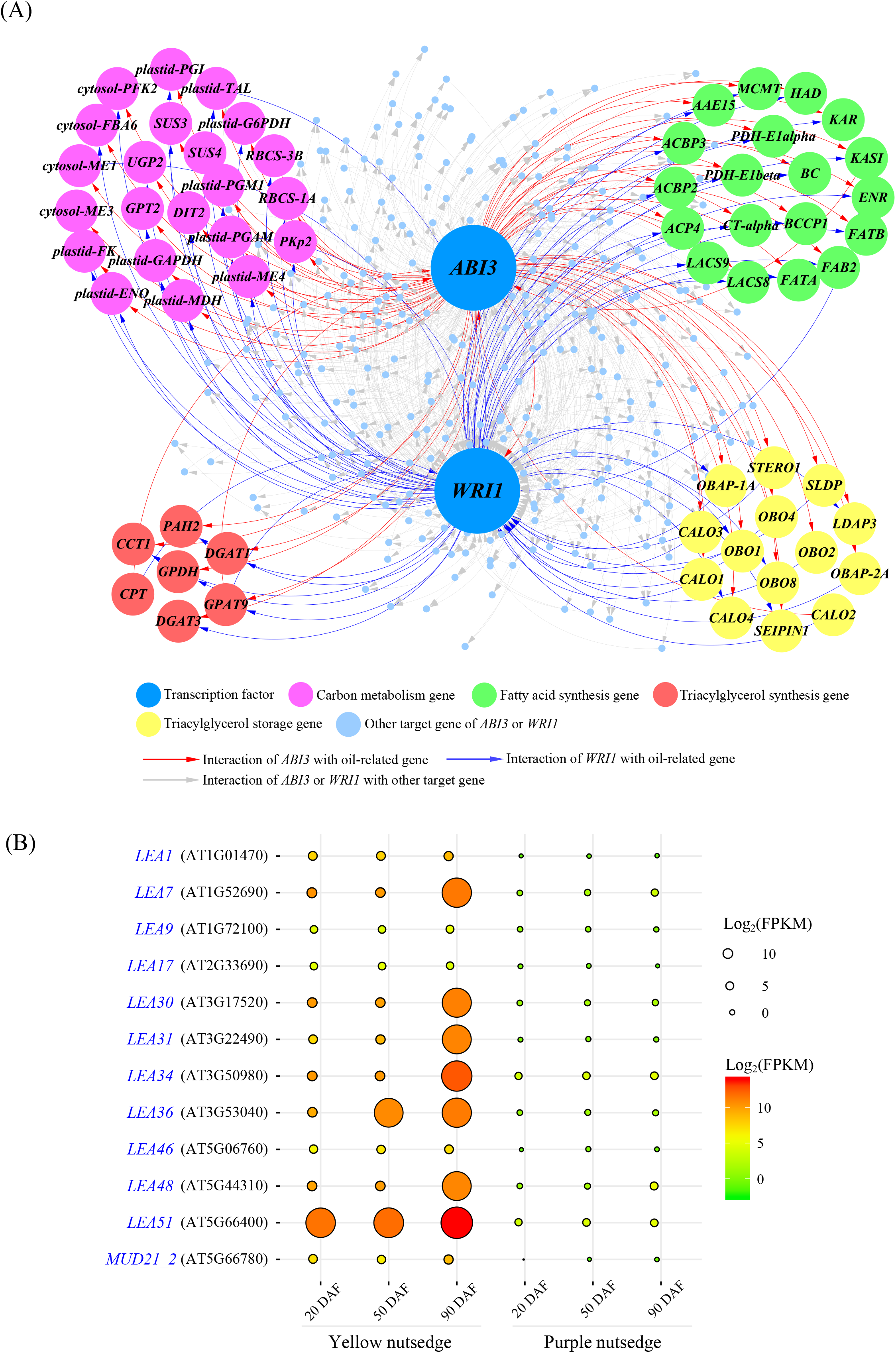
Co-expression network for genes interacting with *ABI3* and *WRI1*. (A) Gene co-expression network of *ABI3* and *WRI1* with selected putative target genes. Genes are represented as nodes and interacting connections are represented as edges that model significant correlation. (B) Balloon plot showing temporal changes in transcript levels (as log2(FPKM)) for genes encoding for late embryogenesis abundant (LEA) proteins. DAF, days after tuber formation.

It was noted that a large number of genes encoding for late embryogenesis abundant (LEA) proteins, which were charactered well as under control of ABI3 (Parcy et al, 1994; Delahaie et al, 2013; González-Morales et al, 2016), are also identified as potential targets of ABI3 and WRI1, as shown in the co-expression network (Supplemental Table S4). Much higher transcripts for these LEA proteins in yellow nutsedge than in purple nutsedge reflect the enhanced desiccation tolerance of yellow nutsedge (Fig. 10B), which allow its tubers can be stored for long time (Turesson et al, 2010).

Collectively, these results reinforce the fact that WRI1 and ABI3, as the master regulators of oil accumulation, do not only regulate gene expressions involved in fatty acid synthesis, but also control the expression of genes encoding for seed-specific oil-body storage proteins (Mönke et al, 2012; Ischebeck et al, 2020; Pouvreau et al, 2020; Tian et al, 2020). Therefore, WRI1-related regulation network of oil production in yellow nutsedge is most likely to differ either from oil seeds or oil fruits, possibly tuber-specific.

## Discussion

In stark contrast to its close relatives of the same Cyperaceae family such as purple nutsedge, Australian bush onion (*Cyperus bulbosus*), priprioca (*Cyperus articulatus*), water chestnut (*Eleocharis dulci*s) that contain exclusively starch or sugar as the major storage reserves in tuberous sink tissues, yellow nutsedge accumulates high amounts of both oil and starch as the main storage compounds in the tubers. This fascinating feature spur us to carry on the present investigation with the aim to elucidate the molecular mechanism and the specific regulatory factors for oil synthesis in yellow nutsedge that distinguishes it from the carbohydrate-rich relatives. In this study, we have systematically carried out a comprehensive comparative analysis of the global expression profiles of genes involved in tuber oil production between yellow and purple nutsedge that lack genome information publicly available. Also, we presented for the first time a public whole transcriptomic profile and framework for purple nutsedge, a traditional medicinal plant used long in India and China.

Our results showed that all the key genes responsible for oil production involved in consecutive pathways of carbon metabolism, FA synthesis, TAG synthesis, and TAG storage are detectable to be expressed and functionally conserved in both yellow and purple nutsedge tubers, indicating that these two nutsedges keep a common set of oil-related genes during their evolution, even though purple nutsedge accumulates low levels of oil in its tuber. Similarly, transcripts for TAG synthesis remained the lowest among the four major metabolic pathways. The transcript patterns with relatively higher expression levels for genes associated with carbon metabolism against FA synthesis were also similar in both species. In addition, the transcript levels for glycolysis genes were much higher in cytosol than in plastid. Moreover, most genes for TAG synthesis were expressed somewhat constantly while for TAG storage were up-regulated during tuber development. All these results point to the fact that tubers of the two species displayed some conserved transcript patterns of oil-related genes, which is similar to non-green heterotrophic oil seeds.

Yellow nutsedge accumulates considerably more oil than purple nutsedge in tubers, suggesting the existence of a tissue-specific transcriptional network controlling oil production. Indeed, differences of expression patterns for oil-related genes were also remarkable between the two species. For example, transcripts for plastidial Rubisco bypass as well as malate and pyruvate metabolism, which are processes destined to pyruvate generation toward for fatty acid synthesis, were much more abundant in yellow nutsedge than in purple nutsedge, suggesting that pyruvate availability in the plastid is an important factor for higher oil accumulation in yellow nutsedge. In addition, yellow nutsedge expressed much higher levels of transcripts for almost all fatty acid synthesis enzymes than purple nutsedge did, implying that transcriptional regulation of fatty acid synthesis genes is another important factor related to high oil accumulation in yellow nutsedge. Furthermore, the large difference of transcripts for TAG storage genes between two species reflected that oil body/lipid droplet biogenesis and/or stability play a significant role in producing high levels of oil in yellow nutsedge. Interestingly, the above-mentioned differentially expressed genes were transcribed in tight coordination with both hub genes of *ABI3 and WRI1*, as showed by co-expression network analysis using WGCNA, implying that ABI3 and WRI1 are potential key factors and master regulators for high oil accumulation in yellow nutsedge. Taken together, all these differences might contribute towards more oil is accumulated in yellow nutsedge than in purple nutsedge.

Many studies have previously tried to elucidate the molecular mechanism that governs the difference of oil content between high- and low-oil lines in other plant species such as soybean (Wei et al, 2008), rapeseed (Li et al, 2006), maize (Liu et al, 2009), sunflower (Troncoso-Ponce et al, 2010) and oil palm (Wong et al, 2017). Comparative transcriptomic analyses of oil palm with date palm mesocarps (Bourgis et al, 2011), and of wild type with transgenic potato tubers expressing Arabidopsis *AtWRI1* (Hofvander et al, 2016) have also been taken to disclose the controlling mechanisms responsible for relatively high or low oil accumulation in these nonseed storage tissues. A common conclusion reached from these studies is that the regulation of gene expression involved in fatty acid synthesis is an important contributor and required for enhanced oil accumulation in diverse plants and tissues. In contrast, our result implied that the main point of controlling oil accumulation in oil tuber may well happen not only at the level of fatty acid synthesis, but also at the level of package and stabilization of fatty acid in the form of TAG. In other words, the flux of fatty acids to oil body formation may ultimately affect tuber oil content.

## Conclusions

A transcriptomic comparison made in this study for the first time revealed several obviously differential transcript patterns of oil-related genes that distinguish yellow nutsedge from purple nutsedge. Higher oil accumulation in yellow nutsedge against purple nutsedge is strongly correlated with upregulation transcripts coding for almost all fatty acid synthesis enzymes and oil body/lipid droplet proteins, specific plastid Rubisco bypass, malate and pyruvate metabolism enzymes, and ABI3- and WRI1-like transcription factors. The most difference is notable for transcripts involved in oil body-related genes between yellow and purple nutsedge. These distinctively expressed genes reflect the differential carbon flux toward oil production between the two nutsedges and may be the key drivers for the high oil accumulation in yellow nutsedge tubers. Together, our study represents an important step toward determining the transcriptional control of key genes responsible for the different oil content between two nutsedge species and provide a useful reference to explore underlying mechanism leading to high oil production in other oil yielding plant species. In addition, this study provides a molecular basis for metabolic engineering or genetic breeding in the future to enhance oil content in yellow nutsedge, or other root or tuber crops through manipulation of the key genes, particularly those related to TAG storage and the master regulator ABI3 as new targets.

## Acknowledgements

This work was supported by National Natural Science Foundation of China (No. 31371692) and Natural Science Foundation of Beijing Municipality (No. 5151001).

## Consent for publication

Not applicable.

## Ethics approval and consent to participate

Not applicable.

## Supplemental material

**Supplemental Table S1**. Summary of transcriptome datasets, assembly and annotation from tuber samples of yellow and purple nutsedge.

**Supplemental Table S2**. Annotation and expression levels for genes associated with carbohydrate metabolism.

**Supplemental Table S3**. Annotation and expression levels for genes associated with lipid metabolism.

**Supplemental Table S4**. Selected genes in tight transcriptional coordination with ABI3 and WRI1.

## Data Deposition

RNA-seq data are available on the National Center for Biotechnology Information (NCBI) Sequence Read Archive (SRA) BioProject under accession number PRJNA671562.

